# Mechanism of ATP hydrolysis dependent rotation of ATP synthases

**DOI:** 10.1101/2022.12.23.521728

**Authors:** Atsuki Nakano, Jun-ichi Kishikawa, Kaoru Mitsuoka, Ken Yokoyama

## Abstract

F_1_ domain of ATP synthase is a rotary ATPase complex in which rotation of central γ-subunit proceeds in 120° steps against a surrounding α_3_β_3_ fueled by ATP hydrolysis. How the ATP hydrolysis reactions occurring in three catalytic αβ dimers are coupled to mechanical rotation is a key outstanding question. Here we describe catalytic intermediates of the F_1_ domain during ATP mediated rotation captured using cryo-EM. The structures reveal that three catalytic events and the first 80° rotation occur simultaneously in F_1_ domain when nucleotides are bound at all the three catalytic αβ dimers. The remaining 40° rotation of the complete 120° step is driven by completion of ATP hydrolysis at α_D_β_D_, and proceeds through three sub-steps (*83°*, *91°*, *101°*, and *120°*) with three associated conformational intermediates. All sub-steps except for one between *91°* and *101°* associated with phosphate release, occur independently of the chemical cycle, suggesting that the 40° rotation is largely driven by release of intramolecular strain accumulated by the 80° rotation. Together with our previous results, these findings provide the molecular basis of ATP driven rotation of ATP synthases.

## Main

The majority of ATP, the energy currency of life, is synthesized by oxidative phosphorylation, catalysed by ATP synthase and dependent on generation of the proton motive force by respiratory complexes^1–3^. F-type ATP synthases, F_o_F_1_, exist in the inner membrane of mitochondria, the plasma membrane of bacteria, and the thylakoid membrane of chloroplasts, while Archaea and some bacteria express the homologous V-type ATP synthases, called V/A-ATPases^4–8^. F_o_F_1_ consists of a hydrophilic F_1_ domain containing three catalytic sites^9^, and a hydrophobic F_o_ domain housing a proton translocation channel^10,11^ (Fig. 1a). The movement of protons through the translocation channel of F_0_ drives rotation of the γ-stalk and this catalyzes the conversion of ADP to ATP at the α_3_β_3_ dimer catalytic sites referred to as α_E_β_E_, α_T_β_T_, and α_D_β_D_. Upon dissociation from the F_o_ domain, the F_1_ domain can catalyze the reverse reaction, i.e. hydrolysis of ATP driven by rotation of the central γ subunit inside the cylinder comprising α_3_β_3_ (Fig. 1b,c). The catalytic sites are located at interface between the α and β subunit in each catalytic dimer. Most of the residues involved in ATP binding and hydrolysis are found in the β subunit, although residues in the α subunit are also involved^12,13^.

**Figure 1.**
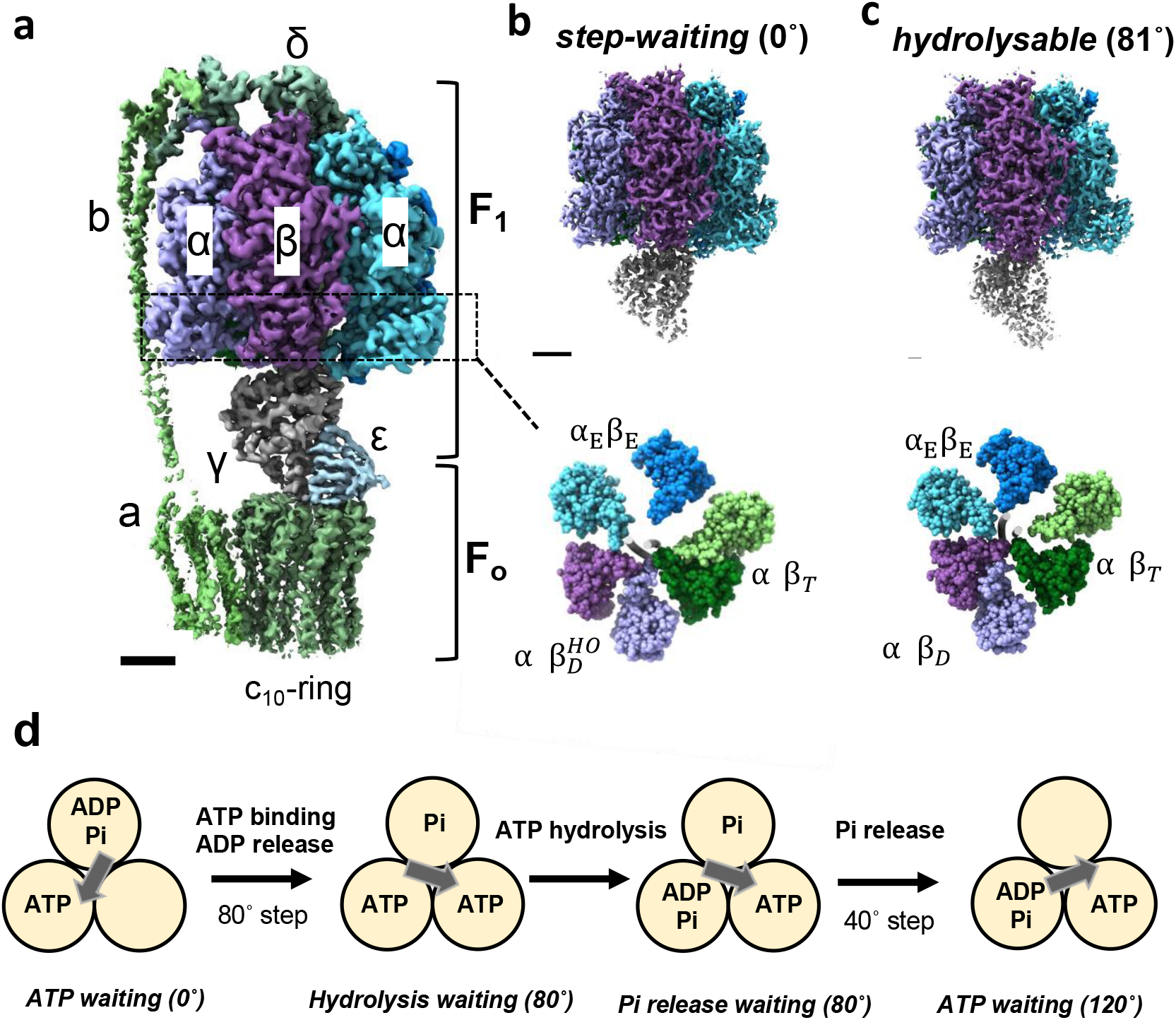
Structures and rotation mechanism of F_o_F_1_. **a** Cryo-EM structure of F_o_F_1_ ATP synthase in the *step-waiting* conformation. Each subunit is colored differently. The F_1_ domain contains three catalytic αβ dimers which surround the γ subunit. **b, c** Side views (*upper*) and bottom section views (*lower*) of the F_1_ domain in the *step-waiting* (0°) and *hydrolysable* (80°) conformational states, respectively. The three catalytic dimers are represented different colors: α_E_ (marine blue) and β_E_ (light blue), α_T_ (moss green) and β_T_ (light green), and α_D_ (purple) and β_D_ (light purple). The β_D_ subunit in *step-waiting* adopts a more open structure, termed as 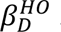. The γ subunits are represented as a grey tube in the center of both α_3_β_3_ sub-complexes. **d** A proposed scheme for chemo-mechanical coupling during a 120° rotation step of the F_1_ domain driven by ATP. In this model, ATP binding to the F_1_ domain immediately initiates the 80° rotation with an associated release of ADP. ATP is hydrolyzed at the 80° dwell position with no associated rotation of γ. It is the release of Pi from the enzyme which is suggested to drive the final 40° rotation.

The binding change mechanism of ATP synthesis at the F_1_ domain by rotation of the γ subunit relative to α_3_β_3_ was firstly proposed by P. Boyer^9,14^. According to this mechanism, the three catalytic dimers adopt different conformational states, and they interconvert sequentially between three the different conformations as catalysis proceeds. Strong support for this asymmetrical F_1_ arrangement was confirmed by the X-ray crystal structure of mitochondrial F_1_ (MF_1_)^13^, which revealed the three catalytic β subunits in different conformational states and with different nucleotide occupancy at the catalytic sites; closed β_DP_ with ADP, closed β_TP_ with ATP analogue, and open β_E_ with no bound nucleotide.

Rotation of F_1_ driven by ATP hydrolysis was directly demonstrated in single molecule observations using bacterial F_1_ from *Geobacillus stearothermophilus^15^*. When using 40 nm gold beads with almost negligible viscous resistance, F_1_ pauses at the 80° dwell position during the 120° rotation step at an ATP concentration close to the *K*m^16^. Histogram analysis of the frequency of dwell times for each dwell position suggests a model in which binding of ATP to F_1_ at 0° causes an initial 80° rotation step of γ subunit, followed by a 40° rotation of the γ subunit due to hydrolysis of ATP and release of phosphate at the 80° dwell position (Fig. 1d)^16,17^. In addition, measurement of the rotational velocity of the rotating probe with viscous resistance or external force suggests that F_1_ converts the hydrolysis energy of ATP into torque with high efficiency^18,19^. However, single-molecule observation experiments only allow observation of the motion of the individual γ subunits to which the observation probe is bound, and do not inform on what events are occurring at each catalytic site.

In order to elucidate the entire F_1_ rotation mechanism driven by ATP, we set out to determine the cryo-EM structures of catalytic intermediates of the F_1_ domain during rotation. By freezing cryo-EM grids at different time points or under different reaction conditions, it is possible to trap intermediate states and thus build up a picture of the chemo-mechanical cycle of motor proteins step by step^20^. Previously, we have determined several intermediate structures of V/A-ATPase revealing an unprecedented level of detail for the ATP hydrolysis cycle in these enzymes. The results showed that catalytic events occur simultaneously at the three catalytic sites of A_3_B_3_ stator, driving the 120° rotation of the central rotor^21^. Sobti *et al*. also performed snapshot analysis using a mutant F_1_ exhibiting slow ATP hydrolysis and obtained structures corresponding to both the 0° and 80° rotation angles of γ subunit^22^. They proposed, however, a different rotation model of rotary ATPase, in which ATP binding and ATP hydrolysis drive the initial 80°rotation and the 40° rotation of F_1_, respectively. As a result, the coupling of the chemo-mechanical cycle of rotary ATPases remains controversial, 60 years after Boyer first predicted the rotary catalytic mechanism of ATP synthase.

Here, we describe multiple reaction intermediates of the wild type of F_o_F_1_ from *G. stearothermophilus* captured using cryoEM. These structural intermediates of the F_1_ domain fill in the missing pieces in the rotation scheme and have allowed us to redraw the complete chemo-mechanical coupling scheme of rotation in F_o_F_1_ powered by ATP hydrolysis.

### Sample preparation and analysis

The ATP hydrolytic activity of *G. stearothermophilus* F_o_F_1_ is very low due to the up-form of the C-terminus of ε subunit^23,24^. In this study, we used the *ΔεCT*-F_o_F_1_ mutant, less susceptible to ε inhibition by truncation of ε-C-terminal helix than the wild-type^24^. The *ΔεCT*-F_o_F_1_ expressed in *E. coli* membranes was solubilized in DDM, then purified as described previously^24^. The *ΔεCT*-F_o_F_1_ exhibited ATPase activity, which obeyed simple Michaelis–Menten kinetics with a *V*_max_ of 164 s^-1^ and a *K*_m_ of 30 μM (Extended Data Fig.1a). The purified *ΔεCT*-F_o_F_1_ was subjected to nucleotide depletion treatment by dialysis against EDTA-phosphate buffer, in order to deplete endogenous nucleotide. The enzymatic properties of the nucleotide depleted enzyme (*ND-ΔεCT*-F_o_F_1_) were almost identical to the non-depleted enzyme (Extended Data Fig.1b). Both depleted and non-depleted Forms of the purified enzymes were concentrated then subjected to a cryo-grid preparation as described below. The *ΔεCT*-F_o_F_1_ is referred to simply as F_o_F_1_, hereafter.

### Structures of *hydrolysable* (81°) and *post-hyd* (83°) at high [ATP]

Cryo-grids were prepared using a reaction mixture of *ND*-F_o_F_1_, containing 26 mM ATP to final concentration, a condition termed high [ATP]. The reaction mixture was incubated for 20 sec at 25°C, and then loaded onto a holey grid, followed by flash freezing. The flow charts showing image acquisition and reconstitution of the 3D structure of F_o_F_1_ at high [ATP] are summarized in Extended Data Fig.2b. The particles containing three rotational states (154 k particles) were subjected to 3D classification focused on the F_1_ domain in order to obtain sub-states of the F_1_ domain. As the result of this classification, three sub-classes of the F_1_ domain were obtained with different γ positions, termed the 80°, 100° and 120° structures (Extended Data Fig.2b). Each structure was further classified by a masked classification.

The two F_1_ domain 80° structures were obtained from 56 k particles using an α_D_β_D_ covering mask (Extended Data Fig.2b). The two 80° structures shared the canonical asymmetric hexamer which adopts the closed α_T_β_T_, closed α_D_β_D_, and open α_E_β_E_ conformation (Fig. 2a, Extended Data Fig.4). The angle of the γ subunits in the two 80° structures relative to the 0° structure were 81° and 83°, respectively (Fig. 3c). The nucleotide bound to the α_D_β_D_ differed between the two structures. In the α_D_β_D_ of the 81° structure, ATP was identified at the catalytic site (Fig. 2c, *left*) and thus we termed the 81° structure as *hydrolysable* given that the density in the corresponding 81° structure at low [ATP] (Fig. 2c, *right*) is likely to contain ADP and Pi (see later for more details). In contrast, the 83° structure obtained at high [ATP] clearly contained ADP and Pi and is designated post-hydrolyzed (*post-hyd*) (Fig. 2c, *center*). The overall structures of α_T_β_T_ and α_E_β_E_ in the *hydrolysable* and *post-hyd* states are mostly identical, but the α_D_β_D_ structures differ slightly. In the *post-hyd*, the *CT* domain of β_D_ was in a slightly more open conformation relative to that in *hydrolysable* (Fig. 3b, Extended Data Fig. 5).

**Figure 2.**
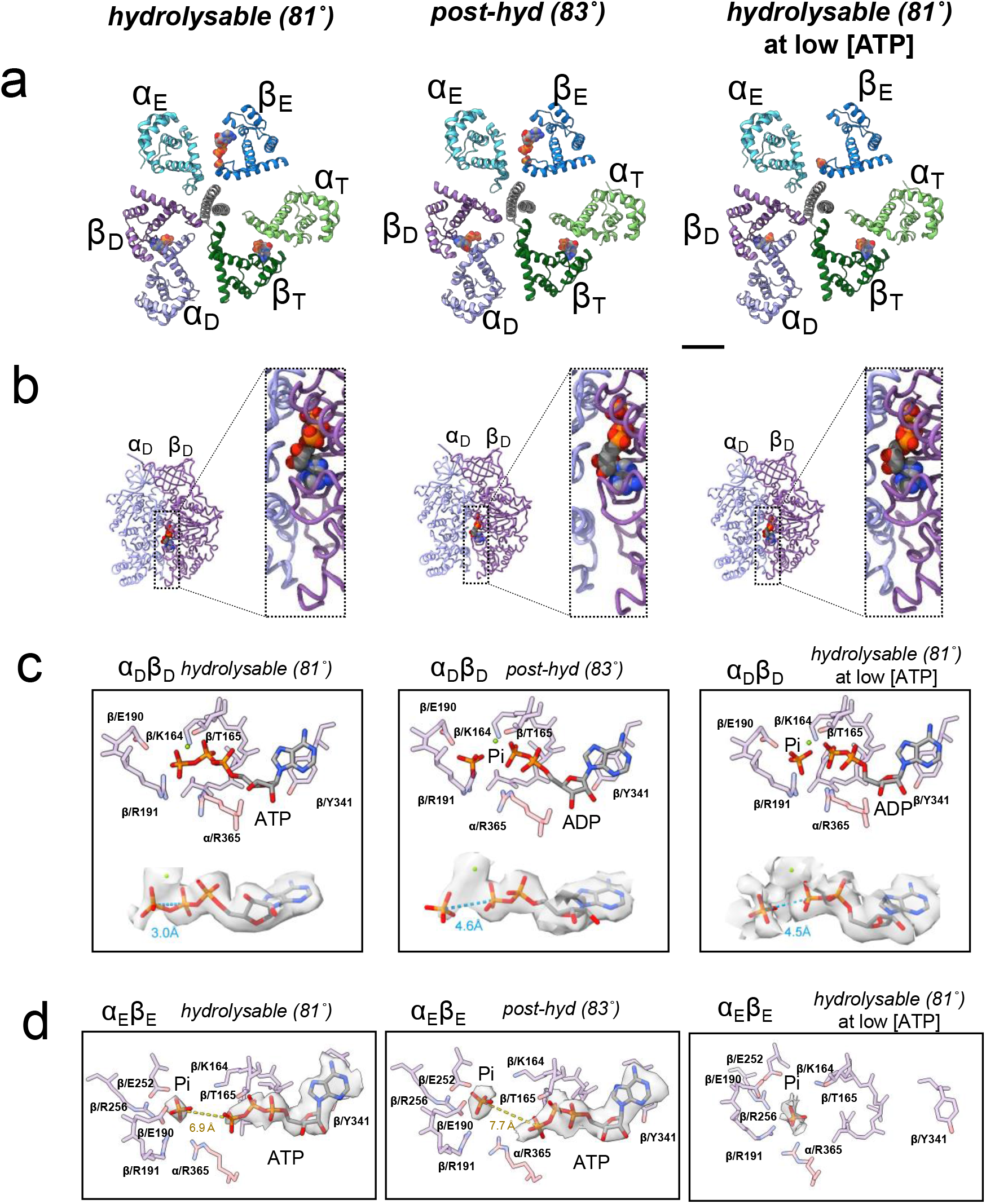
Structure of hydrolysable (*left*) and *post-hyd* (*center*) at high [ATP], and *hydrolysable* (*right*) at low [ATP]. **a** Cross section of the F_1_ domain showing the catalytic sites viewed from the F_o_ side. Each catalytic dimer is shown in ribbon representation and colored as detailed in Figure 1. The bound nucleotides are represented as spheres. **b** Side view structure of α_D_β_D_ dimer in *hydrolysable* at high [ATP] (*left*), post-hyd (*center*), and *hydrolysable* at low [ATP](*right*). The left panel is a magnified view of the catalytic interface at each α_D_β_D_. **c** Structure of the catalytic site in α_D_β_D_. Amino acid residues and bound nucleotide and Pi are represented as sticks. *Lower panels;* Stick representation model of ATP/ADP and Pi with EM density in α_D_β_D_. The distance between γ and β phosphate of the ATP in each conformational state is shown in blue (Å). Each distance was calculated using Chimera software. **d** Structure of the catalytic site in α_E_β_E_. The EM density of ATP / Pi is superimposed onto the model. The distance between Pi and γ phosphate of ATP is shown in yellow (Å).

**Figure 3.**
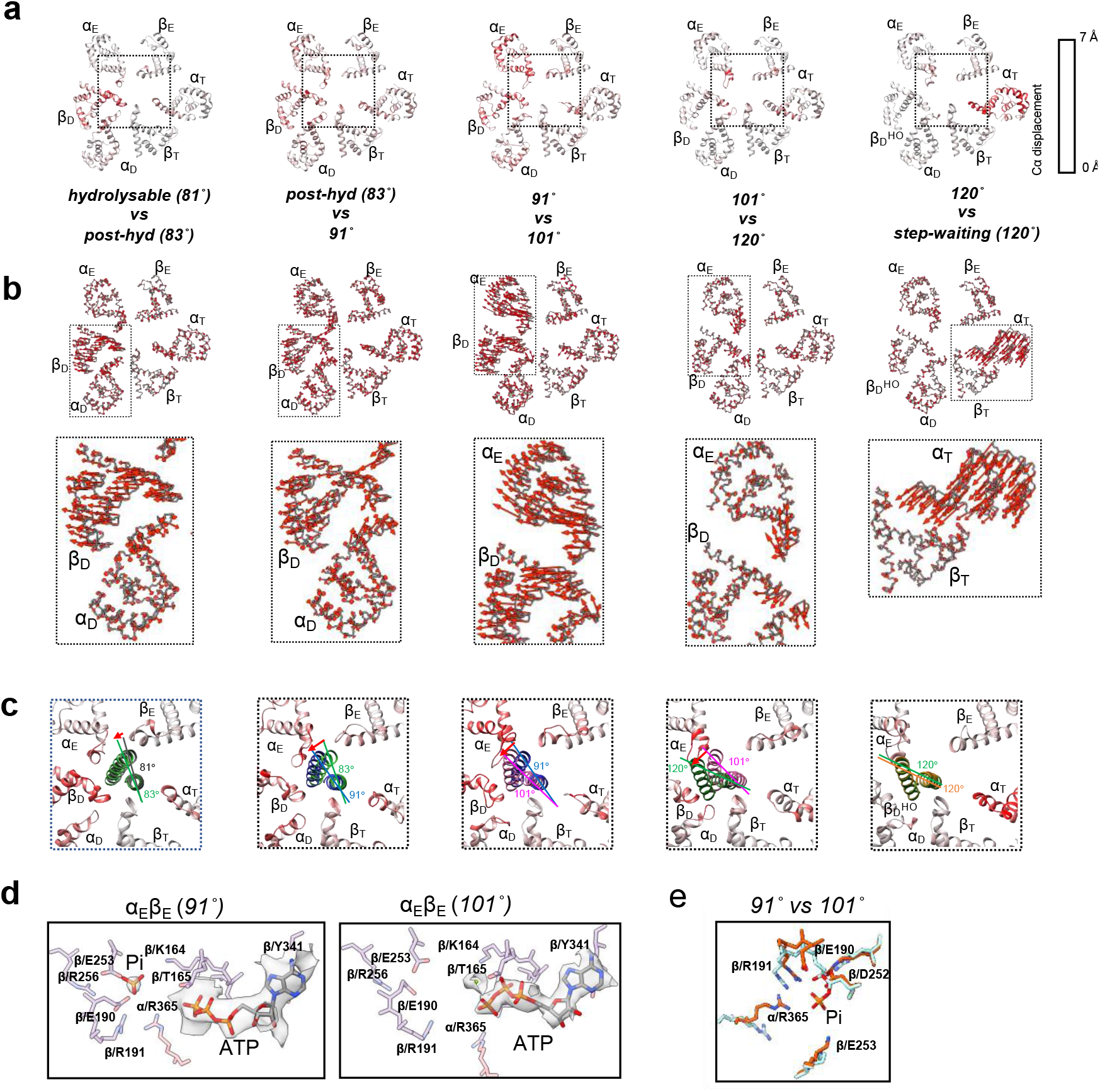
Structure comparison of 6 intermediates captured during the 40° step at high [ATP]. **a** Cross section of the F_1_ domain showing the catalytic site. *Hydrolysable, post-hyd, 91°, 101°*, and *120°* structures are arranged from left to right. The C*α* displacement relative to the next structure (*right side*) is indicated by the red-white color gradient. The dashed square indicates the area of this figure shown in the zoomed in view in **c**. **b** The direction of Cα displacement relative to the next structure is indicated by the red arrow. Longer arrows indicate greater displacement distances. The dashed square indicates the area of this figure shown in the zoomed in view in lower panels. **c** Rotation of the γ subunit and the C-terminal (*CT*) region of the αβ dimer in contact with the γ subunit. The different rotation angles between each γ subunit and that of the next structure are indicated by the different colored lines. The C*α* displacement of *CT* region of the αβ dimer relative to the next structure is indicated by the red-white color gradient. **d** Structures of the catalytic site of 91° and 101° in α_E_β_E_. **e** Superimposition of side chains of α_E_β_E_ in *91°*(*orange*) with those of α_E_β_E_ in *101°* (*light blue*).

In both *hydrolysable* and *post-hyd*, ATP and Pi were identified at the catalytic sites of α_E_β_E_. The Pi is occluded by β/164K, β/190E, β/191K, β/252D, β/256R, and α/365R (Extended Data Fig. 6f). Since β_E_ has an open structure, the site occupied by Pi is separated from the site where ATP is bound. Therefore, in this state the γ-phosphate of ATP does not directly interact with the bound Pi. For other structures obtained under high [ATP], as described in the next section, ATP was identified at the catalytic sites of α_E_β_E_ (Fig. 3d and Extended Data Fig. 4d). This indicates that α_E_β_E_ is capable of binding of ATP to the catalytic site, regardless of the rotary angle of the γ subunit.

### Structures at *91°, 101°* and *120°* rotation angles at high [ATP]

Two additional structures found to be rotated a further 8° and 28° relative to the *post-hyd* (83°), respectively, were classified (Extended Data Fig. 2b). We termed these the *91°* and *101°* structures (Extended Data Fig. 4). In both structures, ATP was identifiable at the catalytic site of α_E_β_E_, but Pi is additionally bound to the *91°* as well as the *hydrolysable* and *post-hyd* forms (Fig. 3d and Extended Data Fig. 4d). In contrast, Pi is not present at the catalytic site of α_E_β_E_ in the *101°* structure, indicating that Pi is released during the 10° rotation of the γ subunit from *91°* to *101°*. The α_E_β_E_ in *101°* adopts a more open structure than that seen in the *91°* structure due to movement of the *CT* domain of α_E_ (Fig. 3b and Extended Data Fig. 4d). This more open arrangement results in a more exposed catalytic site in α_E_β_E_ and thus a decrease in affinity for Pi in the *101° structure*. The geometry of the amino acid residues in β_E_ that coordinate Pi changes due to the release of Pi (Fig. 3e), but no significant change in the conformation of the main chain in β_E_ was observed (Fig. 3a), suggesting that the release of Pi directly does not cause the opening of α_E_β_E_.

Two 120° structures were classified (Extended Data Fig. 2b) according to structural differences in α_T_β_T_. One 120° structure, termed *step-waiting*, has a more closed α_T_β_T_ than the second, termed *120°*. α_T_β_T_ in *120°* is almost identical to that of the *101°* structure (Fig. 3a, b and Extended Data Fig. 7a). The difference in α_T_β_T_ between *120°* and *step-waiting* is the result of movement of the *CT* domain of the α_T_ subunit (Fig. 3a, b and Extended Data Fig. 7). Thus, sequential structural changes occur; a structural change from *101°* to the *120°* associated with the 19° rotation of the γ subunit, followed by closing of α_T_β_T_ which occurs between *120°* and *step-waiting*. Notably, the 19° step of the γ-subunit from the *101°* to the *120°* occurs independently of the catalytic cycle of ATP, such as Pi release and ATP hydrolysis in the F_1_ domain.

Comparing the four conformational states, *hydrolysable, post-hyd, 91°, 101°*, the difference in β_D_ among these structures is more marked than that of β_E_ and β_T_. As the rotational angle of the γ subunit increases, the β_D_ adopts a more open structure due to movement of the *CT* domain but remains bound to both ADP and Pi (Fig. 3b and Extended Data Fig. 5a, b). The motion of the *CT* domain in β_D_ continues until the rotation angle of the γ subunit reaches 101° (Fig. 3b). The α_D_β_D_ dimer in the *101°* and *120°* structures can be described as half open (α_D_β_D_^HO^). This conformational change of α_D_β_D_ also occurs largely independently of the catalytic cycle of ATP, with the exception of the release of Pi occurring between *91°* and *101°*. This suggests that the opening of α_D_β_D_ is largely driven by the release of strain on the molecule accumulated in the initial 80° rotation step.

### Structures of F_1_ domain at low [ATP]

To obtain the structure of the F_1_ domain awaiting ATP binding, F_o_F_1_ containing endogenous nucleotides was mixed with the reaction solution containing 25 μM ATP and 0.2 mg/ml of pyruvate kinase and 4 mM phosphoenolpyruvate in order to regenerate consumed ATP. The solution was incubated at 25°C for 60 seconds and then loaded onto a holey grid followed by flash freezing. The flow charts showing image acquisition and reconstitution of the 3D structure of F_o_F_1_ at low [ATP] are shown in Extended Data Fig. 3b. From the combined 75 k particles for the 80° structure, three intermediate structures, two *post-hyd* and one *hydrolysable* were classified. The two *post-hyd* structures are almost identical, including the angle of the γ subunit, apart from a slightly more open structure of the β_D_ being evident in one of the forms.

Both *hydrolysable* and *post-hyd* structures obtained at low [ATP] were very similar to the corresponding structures obtained at high [ATP] (Extended Data Table 1), but there were significant differences. In the *hydrolysable* and *post-hyd* structures obtained at low [ATP], Pi was identified at the catalytic site of α_E_β_E_, but no ATP density was observed (Figs. 2d and 4a). In addition, ATP molecule was not found in other obtained structures at low [ATP] (Fig. 4a, d and Extended Data Fig. 6d, e) indicating that ATP does not bind to α_E_β_E_ at low [ATP]. This contrasts with the binding of ATP to all 80° structures of α_E_β_E_ obtained at high [ATP] (Extended Data Fig. 4d).

**Figure 4.**
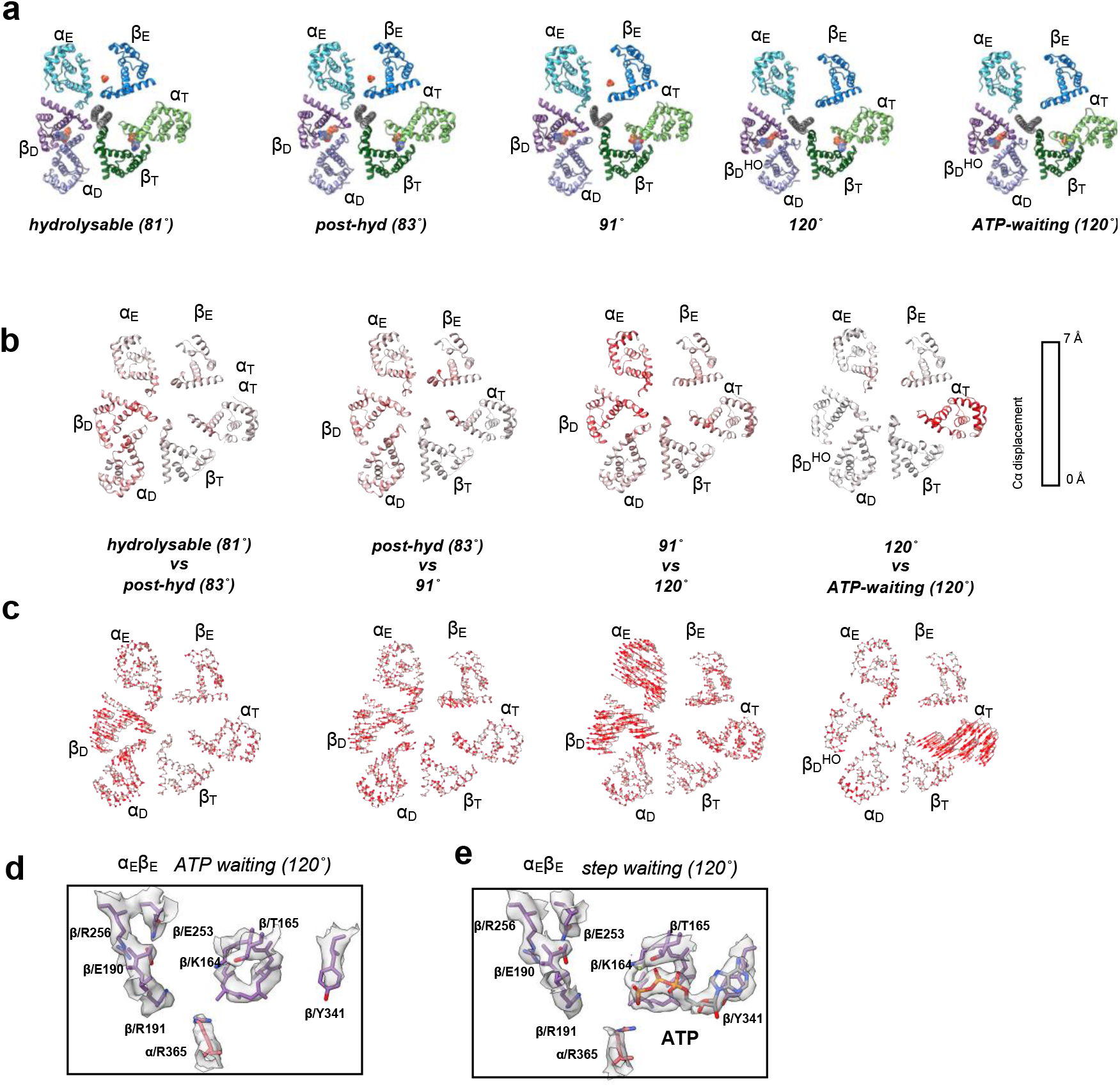
Structure comparison of the 5 intermediates captured at low [ATP]. **a** Cross section of the F_1_ domain showing the catalytic site. Each catalytic dimer is shown in ribbon representation and colored as detailed in Figure 1. The bound nucleotides and Pi are represented as spheres. From left to right, *hydrolysable, post-hyd, 91°* and *120°*, and *ATP waiting* structures are lined up. **b** The C*α* displacement relative to the next structure (*right side*) is indicated by the red-white color gradient. **c** The direction of Cα displacement relative to the next structure is indicated by the red arrow. Longer arrows indicate greater displacement distances. Structures of the catalytic site of *ATP waiting* (**d**) *and step waiting* (**e**) in α_E_β_E_ with EM density. ATP is represented by the orange and blue sticks.

As described above, ADP and Pi were identified at the α_D_β_D_ binding site in *hydrolysable* at low [ATP] (Fig. 2c), indicating that ATP and (ADP + Pi) are in equilibrium at the catalytic site of α_D_β_D_ in *hydrolysable*. This also suggests that ATP hydrolysis at the catalytic site does not directly cause rotation of the γ subunit, consistent with the binding change mechanism where ATP synthesis at the catalytic site is dependent on rotation of γ subunit^1^.

From the combined 332 k particles for 120° structure at low [ATP], three structures of the F_1_ domain, *91°* and *120°* and *step-waiting*, were identified (Extended Data Fig. 3b). The *101°* structure was not captured in this condition. At high [ATP], the F_1_ domain of *120°* and *step-waiting* were composed of the α_D_β_D_^HO^ with ADP and Pi bound, α_T_β_T_ with ATP, and α_E_β_E_ with ATP (Extended Data Fig. 4). However, at low [ATP], ATP nor Pi density was identified at the catalytic site of α_E_β_E_ in either the *120°* and *step-waiting* states (Fig. 4a, d). We refer to the *step-waiting* containing an empty α_E_β_E_ as *ATP-waiting*.

In summary, ATP was not identified at α_E_β_E_ in all six structures, *hydrolysable, post-hyd, 91°, 120°, andATP-waiting*, obtained at low [ATP]. Comparing these results with the nucleotide occupancy of α_E_β_E_ in all structures obtained at high [ATP] indicates that ATP is capable of binding to α_E_β_E_ independent of the rotation angle of γ in the F_1_ domain (Fig. 5).

**Figure 5.**
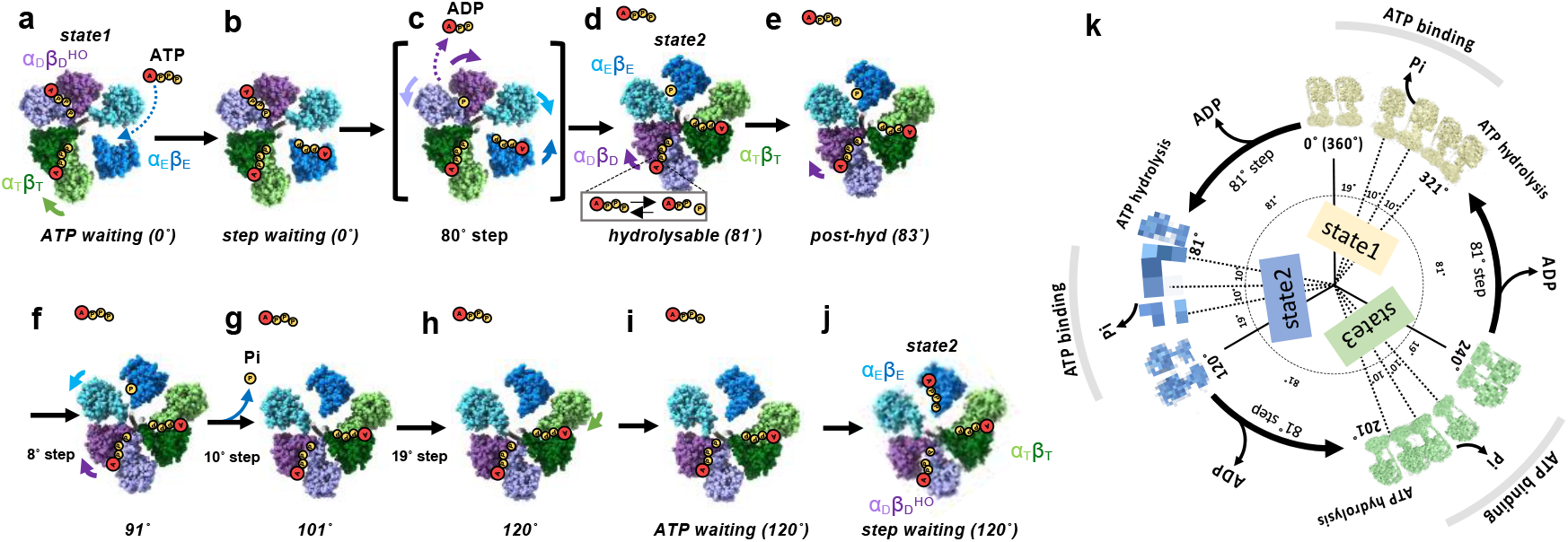
ATP driven rotation scheme of F_o_F_1_. **a** Under low [ATP] conditions, the catalytic site in α_E_β_E_ of *ATP waiting* remains empty. **b** The *step-waiting* is formed by binding of ATP to α_E_β_E_ of *ATP waiting*. **c** The *step-waiting* initiates the 80° rotation step of the γ subunit coupled with structure transition of the three αβ dimers; α_E_β_E_ to α_T_β_T_ with a zippering motion caused by binding of ATP to α_E_β_E_, α_T_β_T_ to α_D_β_D_, and α_D_β_D_^HO^ to α_E_β_E_ with associated release of ADP via an unzippering motion of α_D_β_D_^HO^. **d** ATP bound to α_T_β_T_ is hydrolyzed in the α_D_β_D_ dimer of *hydrolysable* just after the 8O°rotation. **e, f** Once ATP bound to α_D_β_D_ is hydrolyzed, an unzippering motion of α_D_β_D_ (*purple arrows*) proceeds via a 10° rotation step of γ, resulting in *the* structural change of *hydrolysable* to *91°* through *post-hyd*. The outward motion of α_E_ in *91°*(*light blue arrow*) in concert with a further 10° rotation induces release of Pi, resulting in *1O1°* which adopts a more open α_E_β_E_. **g** The final rotation from 101° to 120° occurs without structural change in any of three catalytic dimers. **h** Further motion of the *CT* domain of α_E_ induces the structural between *120°* and, **i** *ATP waiting* without any associated rotation of the γ subunit. **j** The *step-waiting* (120°) is formed by binding of ATP to α_E_β_E_ of *ATP waiting*. The α_E_β_E_ of six intermediates (*hydrolysable, post-hyd, 91°, 101°, 120°, and ATP waiting*) are occupied with ATP at high [ATP], indicating that ATP binds to empty α_E_β_E_ at any rotation angle of γ (*light blue dash arrows*). **k** 360° rotation of F_o_F_1_. CryoEM maps of F_o_F_1_ obtained at high [ATP] are placed in a circular arrangement according to the angle of the γ subunit. The three state maps are represented by yellow (state1), blue (state2), and green (state3), respectively.

### Rotation scheme of F_o_F_1_

In this study, we have captured multiple structures of F_1_ domain and from these we can reveal the rotation scheme that occurs during the full 120° rotation (Figs. 5 a-j, Supplementary movie 1). The first 80°rotation occurring via *step-waiting* of rotational state 1 is coupled with structural transition of three catalytic dimers, α_E_β_E_ to α_T_β_T_, α_T_β_T_ to α_D_β_D_, and α_D_β_D_^HO^ to α_E_β_E_ (Fig 5b, c). The resultant *hydrolysable* is in rotational state 2 (Fig. 5d). We further classified the three rotational states of F_o_F_1_ from the single-particle images of each F_1_ domain structure (Extended Data Figs. 2b and 3b), enabling us to reproduce the 360° rotation scheme of F_o_F_1_ upon hydrolysis of the three ATPs (Fig. 5k). Comparison and interpretation of these intermediate structures provides us with several important insights into the coupling of the chemo-mechanical cycle of ATP hydrolysis driven rotation of F_o_F_1_.

### Tri-site mechanism

ATP binds to empty α_E_β_E_ of the F_1_ domain at any rotation angle, as shown in Fig. 5. Based on single molecule observation experiments for F_1_-ATPase, the coupling scheme for chemo-mechanical cycle of F_1_-ATPase proposed that rotation occurs simultaneously with ATP binding to the structure at a rotation angle of 0° (Fig. 1d)^16–18^. In contrast, structural snapshots of V-type ATP synthase (V/A-ATPase) revealed that the 120° rotations occur after all three catalytic sites are occupied by nucleotides (Extended Data Fig. 8a)^21^. The snapshot analysis of F_o_F_1_ structures in this study indicates that binding of ATP to empty α_E_β_E_ in *ATP-waiting* results in *step-waiting* in which all three catalytic sites are occupied by nucleotides, then the first 80°rotation starts (Fig. 5c). This means that ATP binding to the F_1_ domain does not immediately cause the 80° rotation. Our results clearly demonstrate that F_o_F_1_ and V/A-ATPase share a common tri-site mechanism in which nucleotide occupancy transitions between 2 and 3 binding sites during continuous catalysis^2^.

### The 80° dwell

A further major insight is that the 80° dwell during the 120° step observed in F_1_-ATPase is the result of waiting for ATP hydrolysis at the α_D_β_D_. Either ATP or ADP and Pi was observed at the α_D_β_D_ in *hydrolysable* (Fig. 2c), indicating that hydrolysis of ATP proceeds at α_D_β_D_ in *hydrolysable*. Notably, the α_D_β_D_ in *hydrolysable* has the most closed conformation, adopting a slightly more open conformation in the following *post-hyd* state, (Fig. 2b and Extended Data Figs. 4c and 5a, b). In the α_D_β_D_ of *hydrolysable*, the γ-phosphate of ATP interacts with β/Glu190 and β/Arg192 residues, and the adenine ring interacts with the β/Tyr341 in the *CT* domain (Fig. 2c). Therefore, ATP bound to the catalytic site of α_D_β_D_ stabilizes the closed conformation of the β subunit. Once ATP at the catalytic site is hydrolyzed, α_D_β_D_ is then able to structurally transition to a more open conformation (Extended data Figs. 5c, d). Indeed, the structure of α_D_β_D_ gradually becomes more open as the rotation of the γ subunit proceeds, to the half open α_D_β_D_^HO^ state visible in the *120°* ATP/*step waiting* state (Fig. 5e-i). Based on the dwell time analysis at the 80°rotation angle of F_1_, it has been proposed that two or more catalytic events, including ATP hydrolysis and Pi release, contribute to the 80° dwell^16,17^. Our results support that the dwell at this rotation angle is due mainly to ATP hydrolysis, but is also dependent on the structural transition from the *hydrolysable* to the *post-hyd states* (Figs. 5d, e).

### The subsequent 40° rotation step

The third finding is that the final 40°rotation of the 120°step is composed of three short sub-steps (Figs. 5e-h), that are largely uncoupled from ATP hydrolysis occurring in the F_1_ domain. In previously suggested rotational model of F_1_, the dissociation of Pi at 80° is reported to drive the last 40° of rotation (Fig. 1d) ^17,25^. However our structures reveal that the Pi remains both following rotation beyond 80°, and indeed the structure of α_E_β_E_ without Pi in *101°* indicates that release of Pi occurs during the structural transition from *91°* to *101°*. The rotation of the γ-subunit associated with Pi release is only 10° (Figs. 5f, g), suggesting that the contribution of conformational changes to Pi release (or Pi release by conformational change) is small. Other conformational changes, such as the 8° step of *post-hyd* to *91°* and the 19° step between *101° and 120°*, occur independently of the ATP hydrolysis cycle. In addition, the α_T_β_T_ of *ATP-waiting* adopts a more closed conformation after the structural transition from *120°* to *step-waiting* without rotation of the γ subunit (Fig. 3b, *right*). The only explanation for these spontaneous conformational rearrangement, uncoupled from chemical reactions, is that they are caused by the release of strain inside the molecule. The 80° and 120° (0°) structures identified in the F_1_ domain of F_o_F_1_ are also observed in F_1_-ATPase^22^. Therefore, the strain inside the molecule that drives the 40° step is most likely of F_1_ origin. For instance, a comparison of the γ subunit in the 101° and 120° structures shows a slight structural difference (Extended Data Fig. 8). Together, our results strongly suggest that these structural changes, including the 10° step of *91°* associated with Pi release, is driven by the release of the internal molecular strain accumulated by the initial 80° rotation.

### Principle of rotation mechanism of ATP synthases

Taken together, these structural snapshots of the F_1_ domain indicate that F_o_F_1_ functions via a similar mechanism to the ATP-driven rotation of V/A-ATPase that we have previously demonstrated^21^ (Extended Data Fig. 9a). The binding of ATP to empty AB_open_, corresponding to α_E_β_E_, in V_2nuc_ with two nucleotides bound Forms V_3nuc_ with nucleotides bound to all three AB dimers (AB_open_ with ATP, AB_semi_ with ATP, and AB_closed_ with ADP and Pi). Three distinguishable catalytic events occur at the three AB dimers simultaneously and these events contribute to the first 120° step of the rotor in a concerted manner (Extended Data Fig. 9b)^21^. Considering each 120°step, the rotation mechanism of F_o_F_1_ and V/A-ATPase is almost identical. ATP binds to the F_1_ domain of *ATP-waiting*, resulting in the *step-waiting* where the three catalytic sites are occupied with nucleotides, corresponded to V_3nuc_. *Step-waiting* comprises α_E_β_E_ with ATP, α_T_β_T_ with ATP, and α_D_β_D_^HO^ with ADP and Pi. Each conformational change, from α_E_β_E_ to α_T_β_T_, α_T_β_T_ to α_D_β_D_^HO^, and α_D_β_D_^HO^ to α_E_β_E_ occurs simultaneously, along with the full 120° rotation of the of γ-subunit, and with hydrolysis of ATP in the α_T_β_T_ and release of products (ADP and Pi) from α_D_β_D_^HO^ (Extended Data Fig. 9d). One of the driving events for this structural transition is the conformational change from the ATP-bound α_E_β_E_ to the more closed α_T_β_T_, which can be explained by a zipper motion of α_E_β_E_ occurring upon ATP binding^26^. Our structural analysis indicates that the ATP bound to α_T_β_T_ is hydrolyzed as a result of the conformational change from α_T_β_T_ to α_D_β_D_ which occurs just after the 80° rotation angle (81° to 83°) (Fig. 5). This subsequently leads to the conformational change of α_T_β_T_ to α_D_β_D_^HO^ that occurs during the further rotation to 120° (Extended Data Fig. 9d). The conformational change from α_T_β_T_ to α_D_β_D_^HO^ is likely to occur spontaneously since it involves ATP hydrolysis, an exergonic reaction. The full 120°rotation of the γ-subunit, coupled with spontaneous reactions occurring at both α_E_β_E_ with ATP and α_T_β_T_ with ATP, reduces the affinity For ADP and Pi at α_D_β_D_^HO^, resulting in the release of ADP during the first 80° rotation step and Pi during the following 40°rotation step.

## Discussion

The proposed mechanism of rotation of F_o_F_1_ in this study differs considerably from the model proposed by previous cryo-EM structural snapshots of the F_1_-ATPase, where the 40° rotation is driven by hydrolysis of ATP accompanied by simultaneous release of ADP and Pi^22^. This difference may be due to the use of a mutant F_1_-ATPase (β/E190D) that significantly slows ATP hydrolysis at α_D_β_D_. As a result, only the 80° structure with α_D_β_D_ bound to ATP was identified in the previous study, which would have led to the conclusion that the 40° step was driven by ATP hydrolysis.

Taken together, our comprehensive study demonstrates that the F_o_F_1_ ATP synthase uses the chemical energy of ATP hydrolysis to drive the 80° rotation of the γ subunit and this causes the internal molecular strain which drives the last 40° rotation step, resulting in minimal heat dissipation of the chemical energy. As a result, the coupling of the chemo-mechanical cycle in F_1_ domain is achieved at almost 100 % efficiency, as previously demonstrated for F_1_-ATPase by single molecule observation experiments^2,18,19^.

## Methods

### Sample preparation

For *ΔεCT*-F_o_F_1_ ATP synthase from *G. stearothermophilus*, we deleted the C-terminal domain (83-133 amino acids) of ε from the expression vector PTR19-ASDS^27^. The expression plasmid was transformed into *E. coli* strain DK8 in which the endogenous ATP synthase was deleted. Transformed *E. coli* cells were cultured in 2xYT medium for 24 hours. Cultured cells were collected by centrifugation at 5000 x g and suspended in lysis buffer (50 mM Tris-Cl pH 8.0, 5 mM MgCl_2_, and 10% [w/v] glycerol). The membrane fraction was resuspended in solubilization buffer (50 mM Tris-Cl, pH 8.0, 5 mM MgCl_2_, 10% [w/v] glycerol, and 2% *n-dodecyl-D-maltoside* [DDM]), and then ultracentrifuged at 85,000x*g*. The enzyme in the supernatant was affinity purified using a nickel-nitrilotriacetic acid (Ni^2+^-NTA) column. For protein samples used under high ATP conditions, endogenous nucleotides were removed from *ΔεCT*-F_o_F_1_ by dialysis with phosphate EDTA buffer containing 200 mM sodium phosphate, pH 8.0, 10 mM EDTA and 0.03% DDM at 25°C for 24 hours. Samples were concentrated to ~500 μl by an Amicon ultra (100 k cut), and then subjected to gel permeation chromatography using a Superose^™^ 6 Increase column equilibrated with 20 mM Tris-Cl, pH 8.0, 150 mM NaCl, 0.03 %DDM. Peak fractions containing F_o_F_1_ were collected and concentrated to ~300 μl using an Amicon ultra (100 k cut) for grid preparation.

### Measurement of ATPase activity

The ATPase activity of purified F_o_F_1_ was assessed at 25°C using a NADH-coupled assay^28^. The assay mixtures contained 50 mM Tris-HCl (pH 8.0), 100 mM KCl, 5 mM MgCl_2_, 2.5 mM phosphoenolpyruvate (PEP), 100 μg/ml pyruvate kinase (PK), 100 μg/ml lactate dehydrogenase, and 0.2 mM NADH, and a range of concentrations of ATP-Mg. The reaction was initiated by adding 2 pmol of F_o_F_1_ to 2 ml of the reaction mixture. ATPase activity of F_o_F_1_ was measured by monitoring NADH oxidation over time by absorbance at 340 nm.

### CryoEM grid preparation

Holey Quantifoil R1.2/1.3 Mo grids were used for high [ATP] conditions and UltraAuFoil R1.2 and 1.3 grid for low [ATP] conditions were used. Before using holly grids, they were treated to 1-minute glow discharge by the Ion Bombarder (Vacuum Device).

For high [ATP] conditions, 3 μl of 200 mM ATP-Mg was added to 20 μl of 10 μM of enzyme solution, and then the solutions mixed well by pipetting. The mixtures containing 26 mM ATP and 9 μM enzyme were incubated for 20 sec at 23°C. Then, 3μL enzyme solution was loaded onto the grid and blotted for 6 sec with a blot force of 10, drain time of 0.5 sec, and 100% humidity using a FEI Vitrobot (*ThermoFisher*), followed by flash freezing by liquid ethane. For low [ATP] conditions, 2 μl of reaction buffer containing 0.2 M Tris-Cl pH 8.0, 250 μM ATP, 40 mM PEP, 1 M KCl, 5 mg/ml PK was added to 18 μl of 10 μM enzyme solution, and then mixed well by pipetting. The reaction mixtures containing enzyme were incubated at 23°C for 60 sec. Aliquots of 3μL of the reaction mixtures were loaded onto grids and immediately flash frozen as described above. The prepared cryo-grids were stored in liquid nitrogen until use.

### Cryo-EM data acquisition

Cryo-EM imaging was performed using a Titan Krios (FEI/Thermo Fisher Scientific) operating at 300 kV acceleration voltage and equipped with a K3 electron detector (Gatan) in electron counting mode (CDS). SerialEM software was used for data collection. CryoEM movies were collected at a nominal magnification of 88,000 with a pixel size of 0.88 Å/pix. The defocus range was 0.8-2.0 μm, and data were collected at 50 frames. The total electron dose was 60 electrons/Å^2^.

### Image processing

The detailed procedures for single-particle analysis are summarized in Extended Data Figs 2 and 3. RELION 4.0^29^ and CryoSPARC v3.3^30^ were used for image analysis. The conversion of file formats between RELION and CryoSPARC was executed using csparc2star.py in pyem. For both high ATP (5922 movies) and low ATP (8200 movies) conditions, beam-induced drift was corrected using MotionCor2^31^, and CTF estimation was performed using CTFFIND4.1^32^. Particle picking was performed by Topaz^33^. In each dataset, good particles were sorted by 2D classification after template picking from hundreds of images, and 2000-6000 particles were used to train the Topaz model. 1,118,093 high [ATP] particles were and 2,654,860 low [ATP] particles were picked by Topaz and then these were subjected to 2D classification by CryoSPARC, which further selects particles of good quality. Heterogeneous refinement was used to further select particles and classify into different rotation states. The total number of particles selected and classified was 241,668 at high [ATP] and 557,503 at low [ATP] conditions. All particles were combined and a focused 3D refinement was performed on the F_1_ domain. The F_1_ structures with different rotation angles were classified by 3D classification without alignment with masks for the F_1_ domain. Both CTF refinement and Bayesian polishing were carried out multiple times. Finally, we obtained structures with multiple rotational states at 2.6-4.2 Å resolution, estimated by the gold standard Fourier shell correlation (FSC) = 0.143 criterion (Extended Data Fig. 2c and 3c).

### Model building and refinement

The atomic model was built using the previous cryo-EM structure of bacterial F_o_F_1_ PDB *6N2Y*. The β_DP_ has a very different structure in our novel structures, so F_1_ excluding β_DP_ was fitted as a rigid body and β_TP_ was fitted to the β_DP_ density. The initial model was manually modified using COOT and ISOLDE^34^. The manually modified model was refined by the *phenix.real_space_refinement* program^35^. Manual corrections and refinement were iterated until model parameters were improved. The final model was evaluated by MolProbity^36^ and EMRinger^37^.

## Data Availability

Cryo-EM density maps (.mrc files) and atomic models (.pdb files) obtained in this study have been deposited to both the EMDB and PDB. The accession codes (PDBID and EMDBID) are summarized in Extende data Table 2.

## Acknowledgements

We are grateful to all the members of the Yokoyama Lab for their continuous support and technical assistance. Our research was supported by Grant-in-Aid for Scientific Research (JSPS KAKENHI) Grant Numbers 20H03231 to K.Y., 20K06514 to J.K., and Takeda Science foundation funding to K.Y. Our research was also supported by the Platform Project for Supporting Drug Discovery and Life Science Research (Basis for Supporting Innovative Drug Discovery and Life Science Research (BINDS)) from AMED under Grant Number JP17am0101001 (support number 1312), and Grants-in-Aid from the “Nanotechnology Platform” of the Ministry of Education, Culture, Sports, Science and Technology (MEXT). This work was also supported by JST CREST to K.M. (Grant Number. JPMJCR1865).

## Author contributions

K.Y., A.N. designed, performed and analyzed the experiments. A.N. analyzed the data and contributed to preparation of the samples. J.K. and K.M. provided technical support and conceptual advice. K.Y. designed and supervised the experiments and wrote the manuscript. All authors discussed the results and commented on the manuscript. The authors declare no conflicts of interest associated with this manuscript. All data is available in the manuscript or in the supplementary materials.

