## Extended Data for "Mechanism of ATP hydrolysis dependent rotation of ATP synthases"

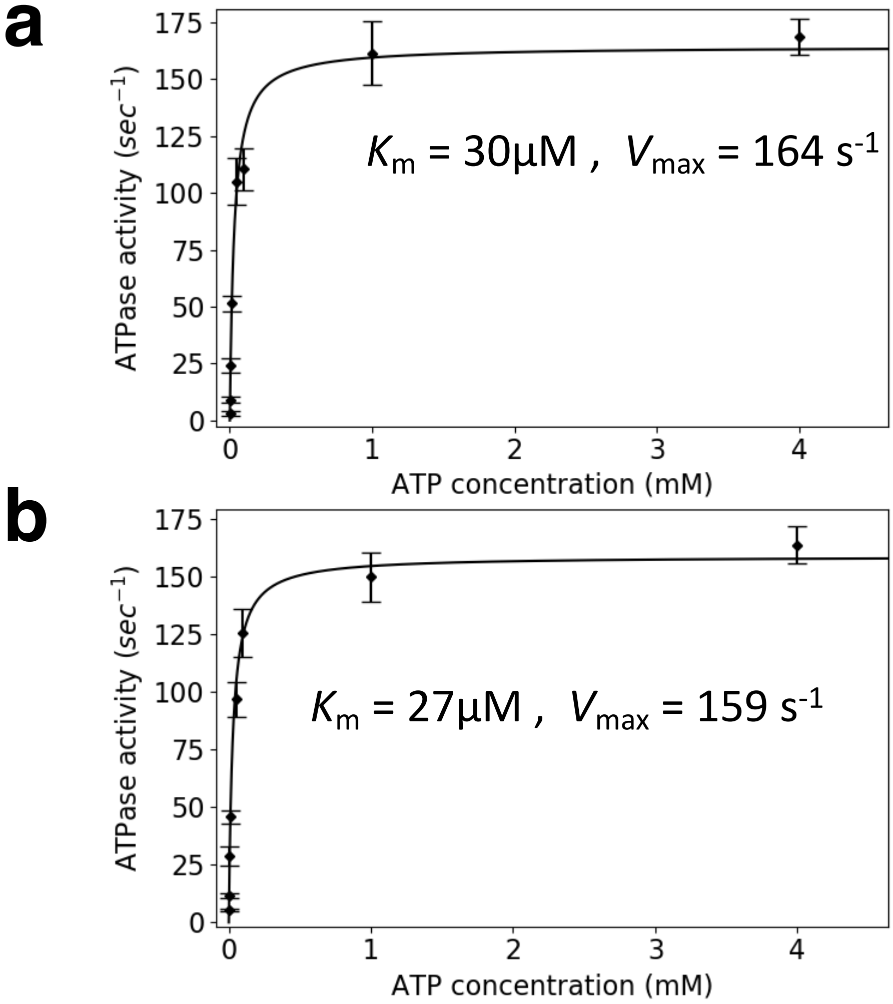


**Extended Data Figure 1.** Enzymatic properties of *ΔεCT*-$F_oF_1$ . **a** Michaelis Menten curves for ATPase activity of ΔεCT-$F_oF_1$ with enzyme kinetics of $K_m$ = 30 $\mu$M and $V_{max}$ = 164 $s^{-1}$, **b** for ATPase activity of the nucleotide depleted enzyme with enzyme kinetics of $K_m$ = 27$\mu$M and $V_{max}$ = 159 $s^{-1}$.

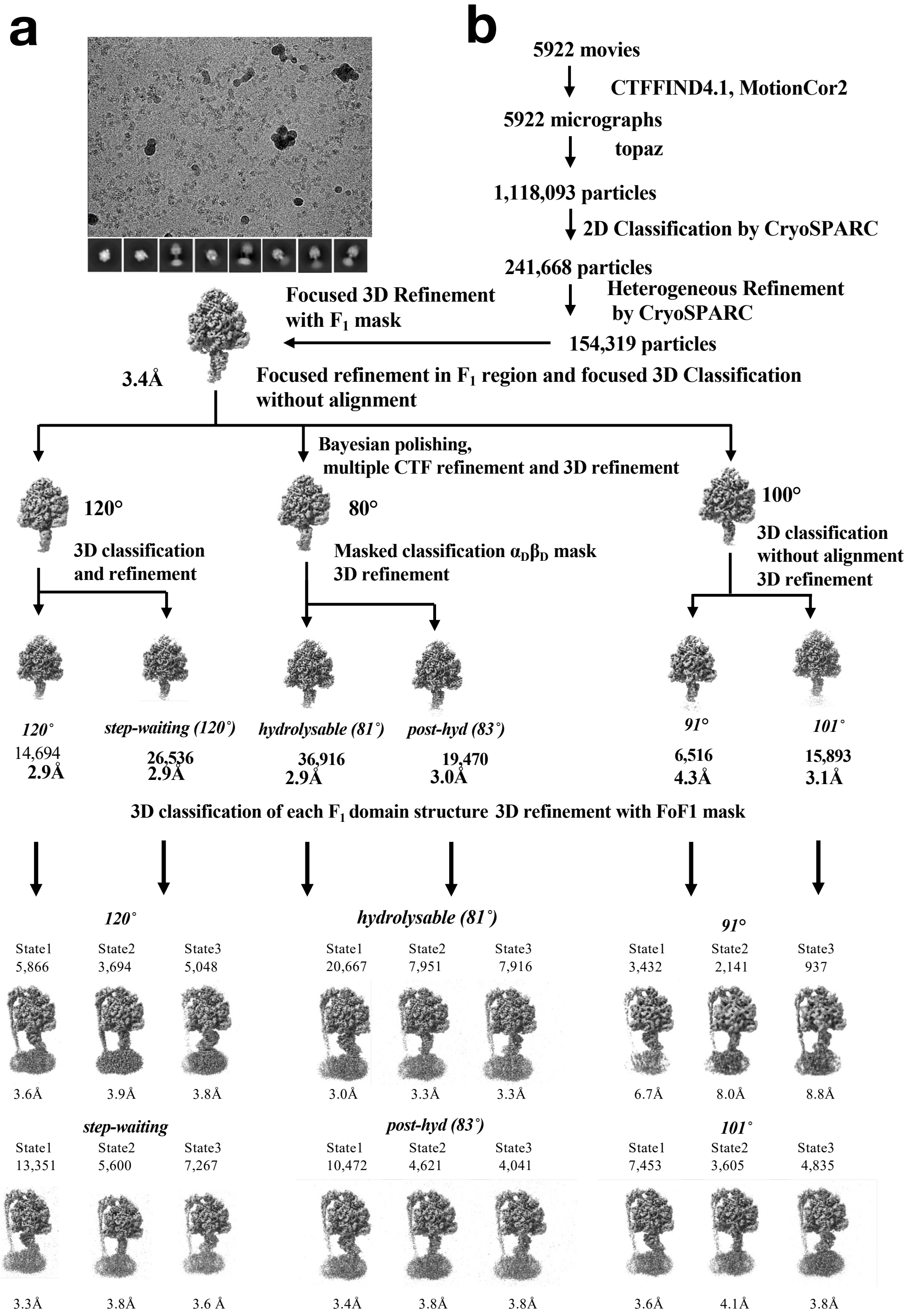

a
b
5922 movies
CTFFIND4.1, MotionCor2
5922 micrographs
topaz
1,118,093 particles
2D Classification by CryoSPARC
241,668 particles
Heterogeneous Refinement by CryoSPARC
154,319 particles
Focused 3D Refinement with $F_1$ mask
3.4Å
Focused refinement in $F_1$ region and focused 3D Classification without alignment
Bayesian polishing, multiple CTF refinement and 3D refinement
120°
3D classification and refinement
80°
Masked classification $\alpha_D\beta_D$ mask 3D refinement
100°
3D classification without alignment 3D refinement
120°
14,694
2.9Å
step-waiting (120°)
26,536
2.9Å
hydrolysable (81°)
36,916
2.9Å
post-hyd (83°)
19,470
3.0Å
91°
6,516
4.3Å
101°
15,893
3.1Å
3D classification of each $F_1$ domain structure 3D refinement with FoF1 mask
120°
State1 5,866 3.6Å
State2 3,694 3.9Å
State3 5,048 3.8Å
hydrolysable (81°)
State1 20,667 3.0Å
State2 7,951 3.3Å
State3 7,916 3.3Å
91°
State1 3,432 6.7Å
State2 2,141 8.0Å
State3 937 8.8Å
step-waiting
State1 13,351 3.3Å
State2 5,600 3.8Å
State3 7,267 3.6 Å
post-hyd (83°)
State1 10,472 3.4Å
State2 4,621 3.8Å
State3 4,041 3.8Å
101°
State1 7,453 3.6Å
State2 3,605 4.1Å
State3 4,835 3.8Å

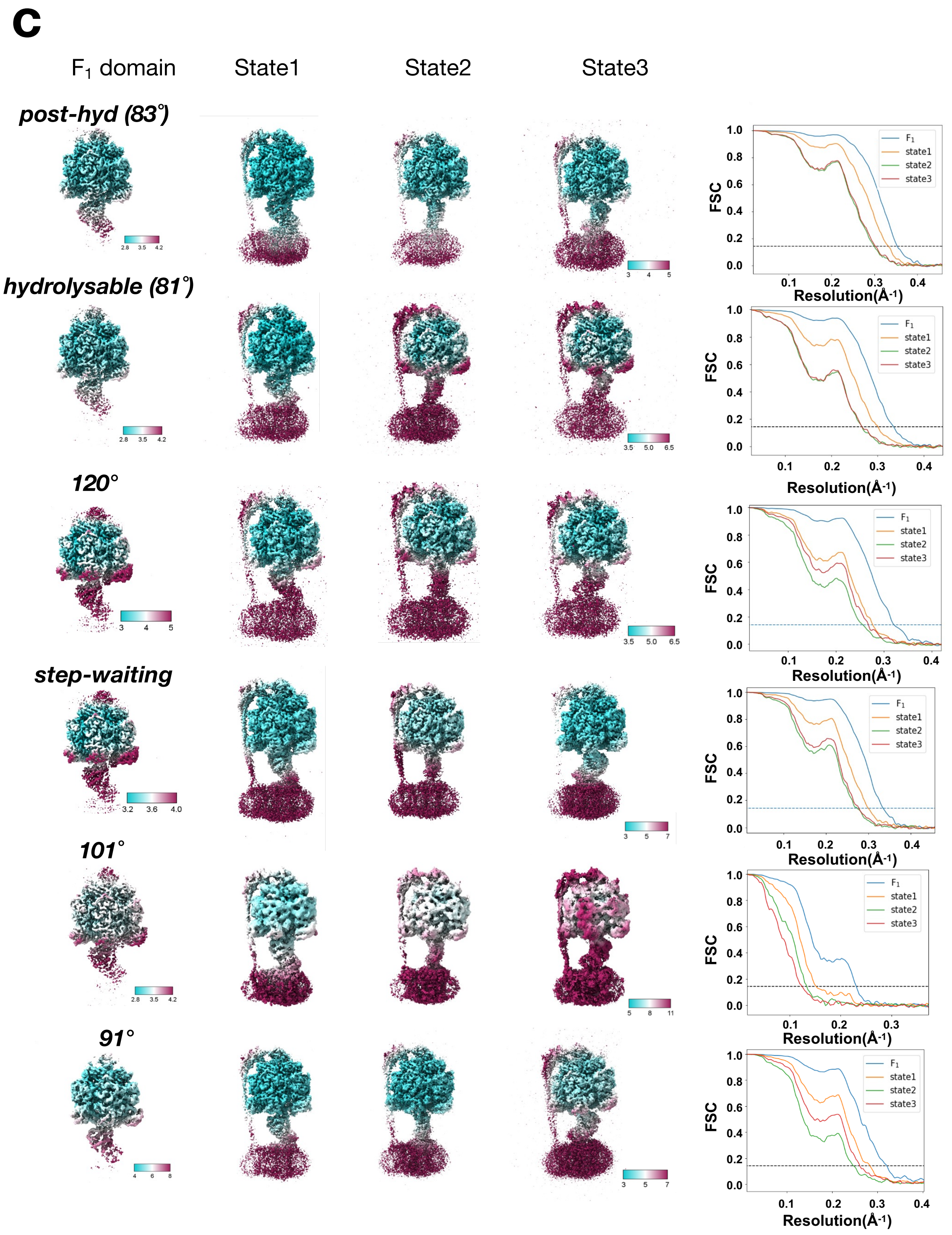


**Extended Data Figure 2** **a** A typical micrograph of *ΔεCT*-$F_oF_1$ at 5mM [ATP](upper) and 2D classes (lower). **b** Flow chart of single particle analysis for *ΔεCT*-$F_oF_1$ at 5mM [ATP]. The selected 154k particles were subjected to focused refinement in the $F_1$ domain and focused 3D classification without alignment, resulting in three classes of 120°, 80°, and 100° rotation of the γ subunit. After 3D classification of the 120° structure, two subclasses were obtained, termed *120°* and *step-waiting*, respectively. *Hydrolyzable* (81°) and *post-hyd* (83°) were identified by $\alpha_D\beta_D$ masked classification of the 80° structure. From the particles of the 100° structure, *91°* and *101°* were obtained by 3D classification without alignment. The six intermediates (*hydrolysable, pot-hyd, 91°, 101°, 120°*, and *step-waiting*) were subjected to further classification with a $F_oF_1$ mask, resulting in three rotational states of six intermediates. **c** Resmaps of six intermediates of the three major rotational states. FSC curves for intermediates of $F_1$ domain and $F_oF_1$ are shown in the right hand panels.

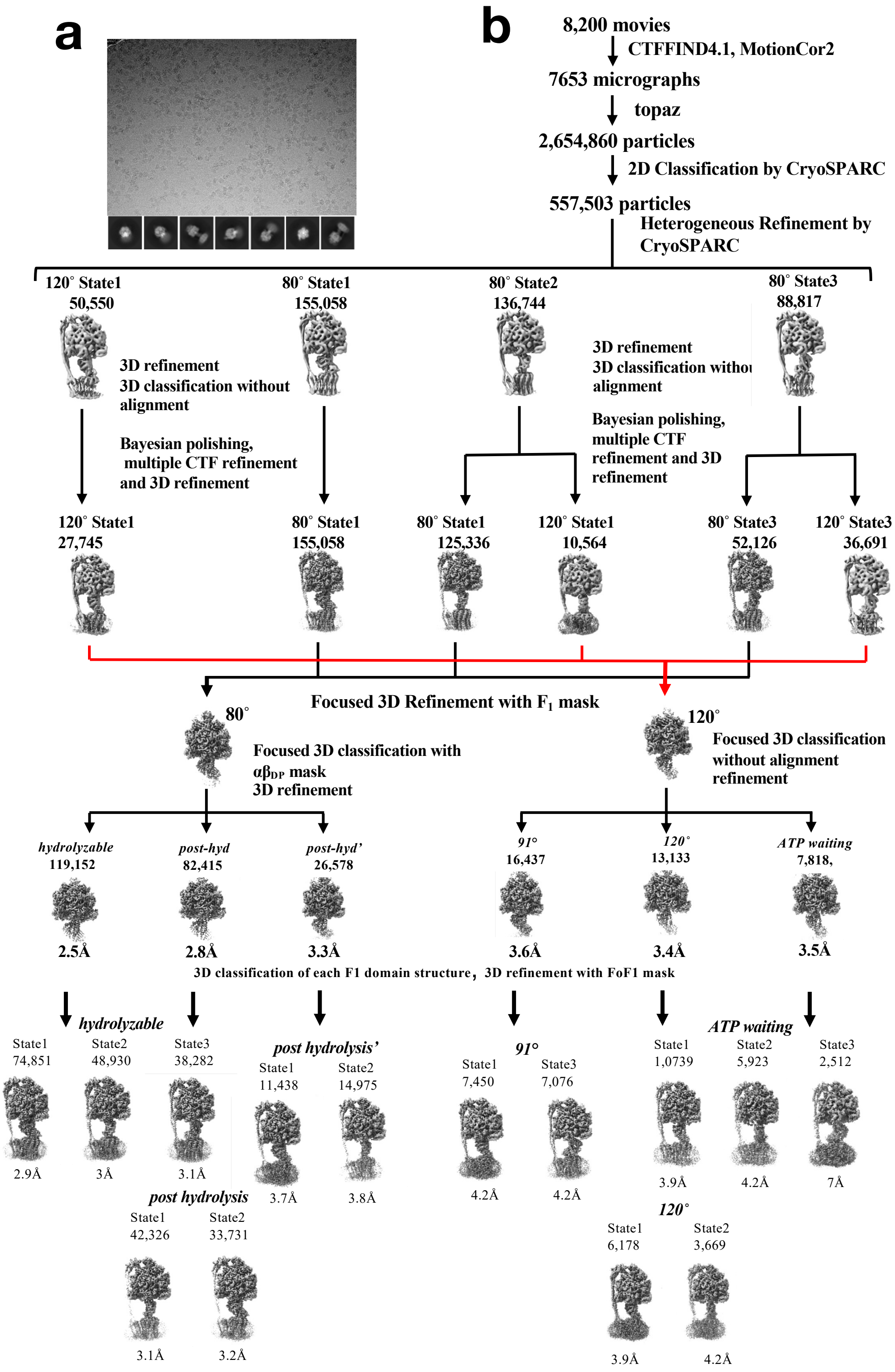
a
b
8,200 movies
CTFFIND4.1, MotionCor2
7653 micrographs
topaz
2,654,860 particles
2D Classification by CryoSPARC
557,503 particles
Heterogeneous Refinement by CryoSPARC
120° State1 50,550
80° State1 155,058
80° State2 136,744
80° State3 88,817
3D refinement
3D classification without alignment
Bayesian polishing, multiple CTF refinement and 3D refinement
120° State1 27,745
80° State1 155,058
80° State1 125,336
120° State1 10,564
80° State3 52,126
120° State3 36,691
Focused 3D Refinement with F1 mask
80°
Focused 3D classification with αβDP mask
3D refinement
120°
Focused 3D classification without alignment refinement
hydrolyzable 119,152
post-hyd 82,415
post-hyd' 26,578
91° 16,437
120° 13,133
ATP waiting 7,818,
2.5Å
2.8Å
3.3Å
3.6Å
3.4Å
3.5Å
3D classification of each F1 domain structure，3D refinement with FoF1 mask
hydrolyzable
State1 74,851
State2 48,930
State3 38,282
2.9Å
3Å
3.1Å
post hydrolysis'
State1 11,438
State2 14,975
3.7Å
3.8Å
post hydrolysis
State1 42,326
State2 33,731
3.1Å
3.2Å
91°
State1 7,450
State3 7,076
4.2Å
4.2Å
ATP waiting
State1 1,0739
State2 5,923
State3 2,512
3.9Å
4.2Å
7Å
120°
State1 6,178
State2 3,669
3.9Å
4.2Å

**c**

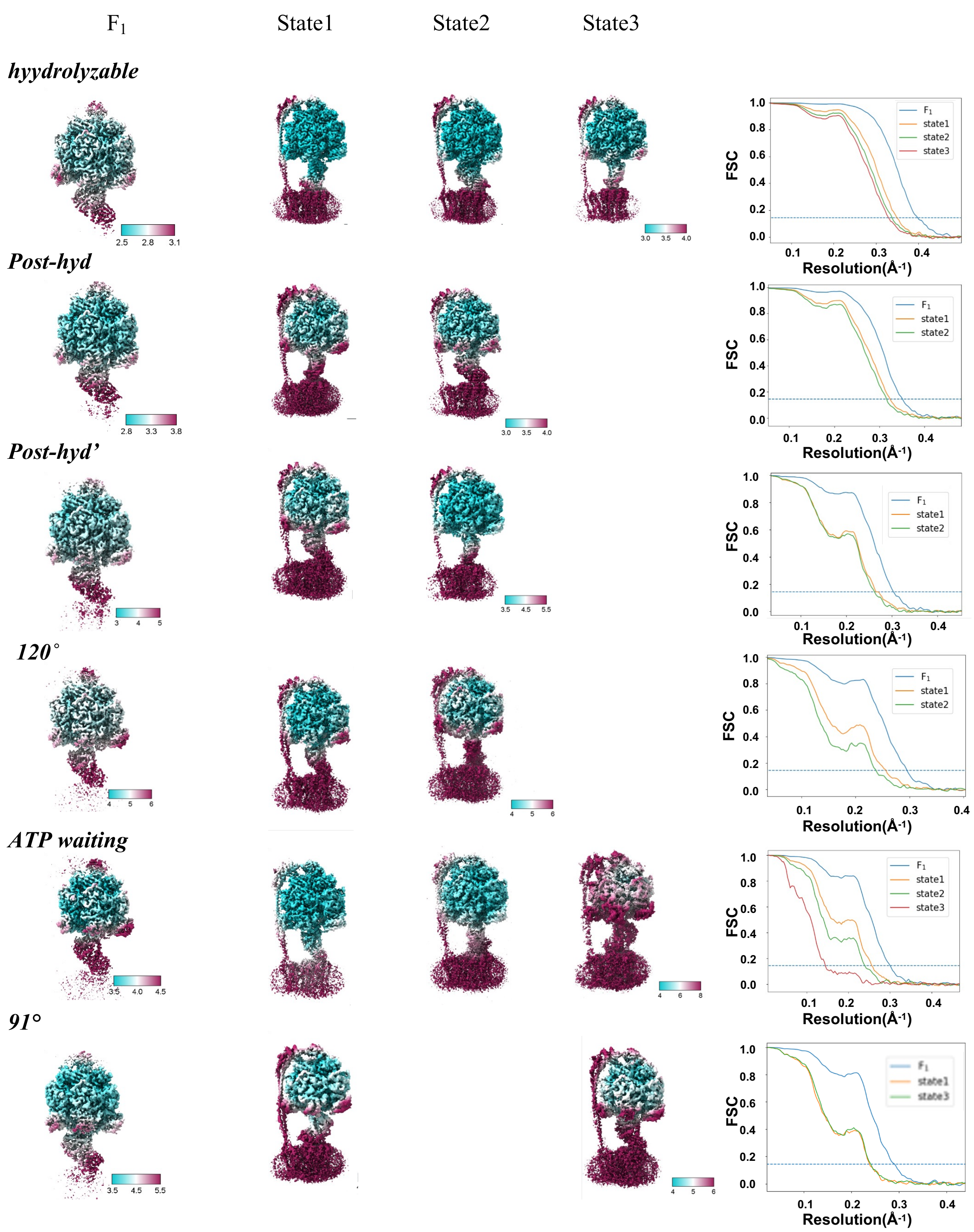


**Extended Data Figure 3.** **a** A typical micrograph of *ΔεCT*-$F_oF_1$ at 50 $\mu$M [ATP](upper) and the 2D classes (lower). **b** Flow chart of single particle analysis for *ΔεCT*-$F_oF_1$ at 50 $\mu$M [ATP]. The Selected 557 k particles were subjected to heterogeneous refinement using CryoSparc. The resulting three 80° structures and one 120° structure were further classified, resulting in three 120° structures and three 80° structures. The particles of 120° and 80° structures were individually combined. The particles of 80° structure were subjected to focused 3D classification using a $\alpha_D\beta_D$ mask, resulting in *hydrolysable* and two *post-hyd* structures. The particles for the 120° structure were subjected to focused 3D classification without alignment, resulting in *91°, 120°*, and *ATP waiting*. The $F_oF_1$ particles of these six intermediates were subjected to further classification using a $F_oF_1$ mask, respectively. Two or three rotational states for six intermediates were classified, respectively. **c** Resmaps of six intermediates of two or three rotational states, respectively. The FSC curves for the intermediates of the $F_1$ domain and $F_oF_1$ are shown in the right hand panels.

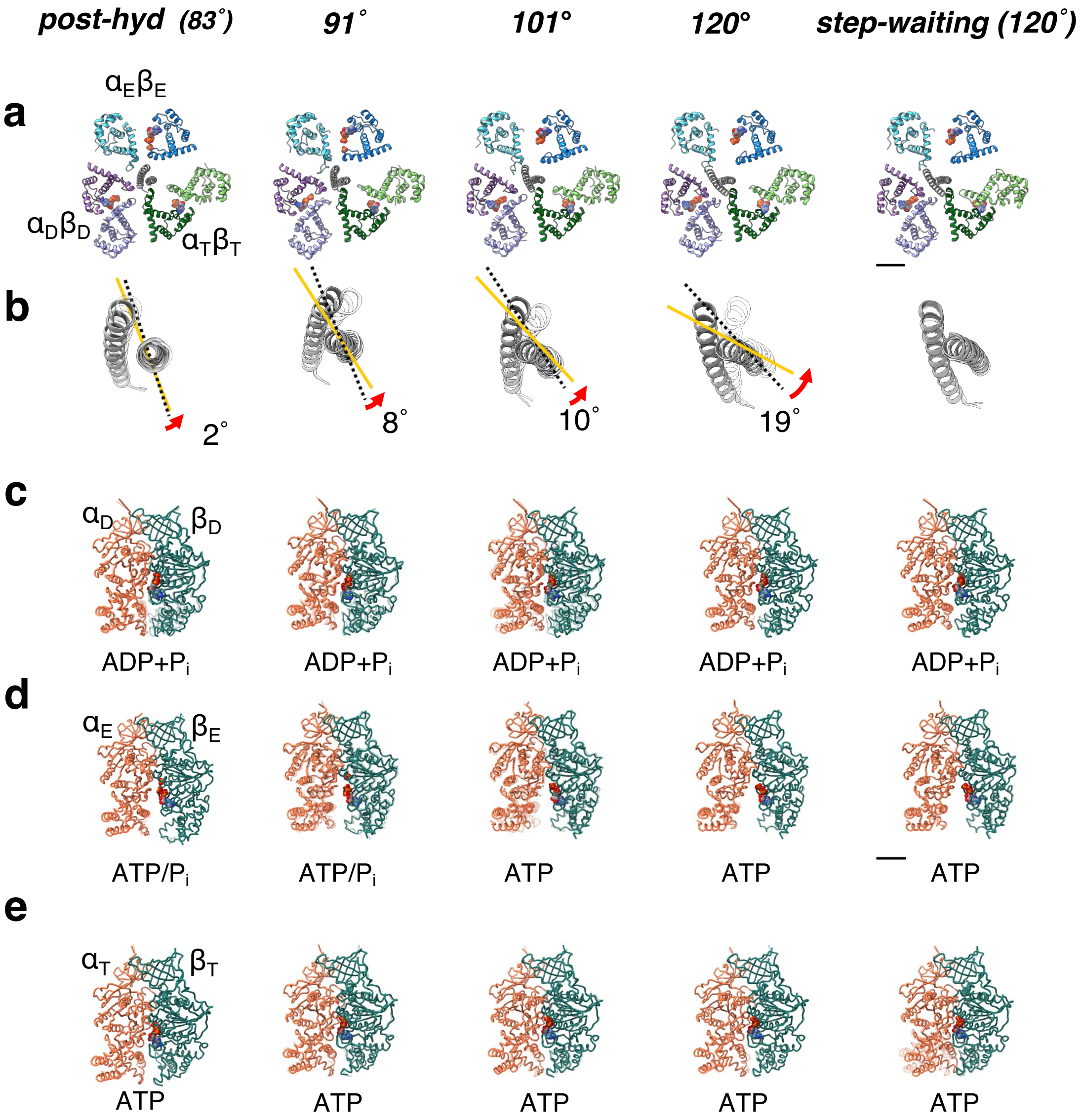


**Extended Data Figure 4. Structure of 5 intermediates captured during the 40° step at high [ATP].** **a** Cross section of $F_1$ domain at the catalytic site. Each catalytic dimer is shown in ribbon representation and colored as detailed in Figure 1. The bound nucleotides are represented as spheres. **b** Rotation angle of the γ subunit relative to that of the 0° structure. The structure of the γ subunit at the previous rotation angle is shown in white. **c-e** Structure of αβ dimers; $\alpha_D\beta_D$ (**c**), $\alpha_E\beta_E$ (**d**), $\alpha_T\beta_T$ (**e**). The bound nucleotide and Pi at the interface of each dimer is represented as spheres with the specific bound molecules labelled under the structures. The superimposed αβ dimers in previous angle are shown translucent, respectively.

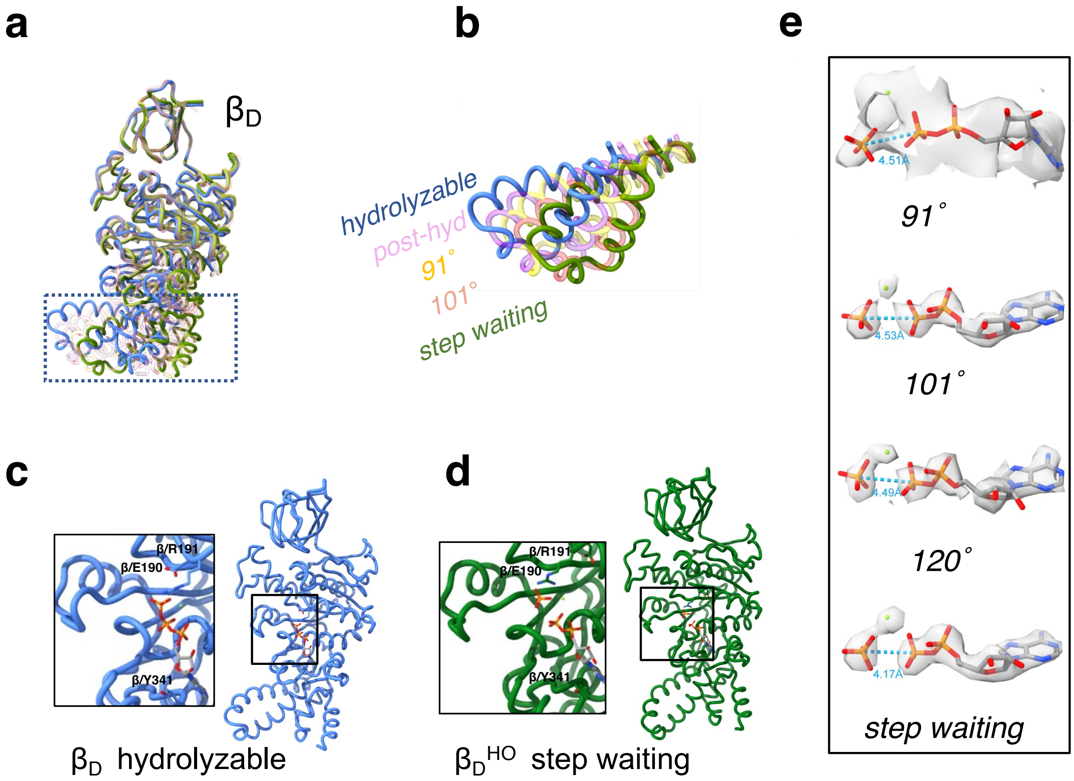


**Extended Data Figure 5.** Unzippering motion of $\alpha_D\beta_D$ upon ATP hydrolysis. **a** superimposition of β subunit in *hydrolysable* with that in *post-hyd, 91°, 101°*, and *ATP waiting*. **b** motion of the *CT* loop helix domain in the conformation changes from *hydrolysable* to *step-waiting*. Structure of $\beta_D$ in *hydrolysable* (**c**) and *step waiting structures* (**d**). The magnified views of the nucleotide binding site are shown in the left panels, respectively. ATP, ADP, Pi and coordinated amino acid residues are shown as sticks. **e** EM density of ATP/ADP+Pi bound to $\alpha_D\beta_D$, with the distance between β-phosphate and γ-phosphate or Pi indicated with a dotted line. Each nucleotide is represented by stick and magnesium ion by green sphere.

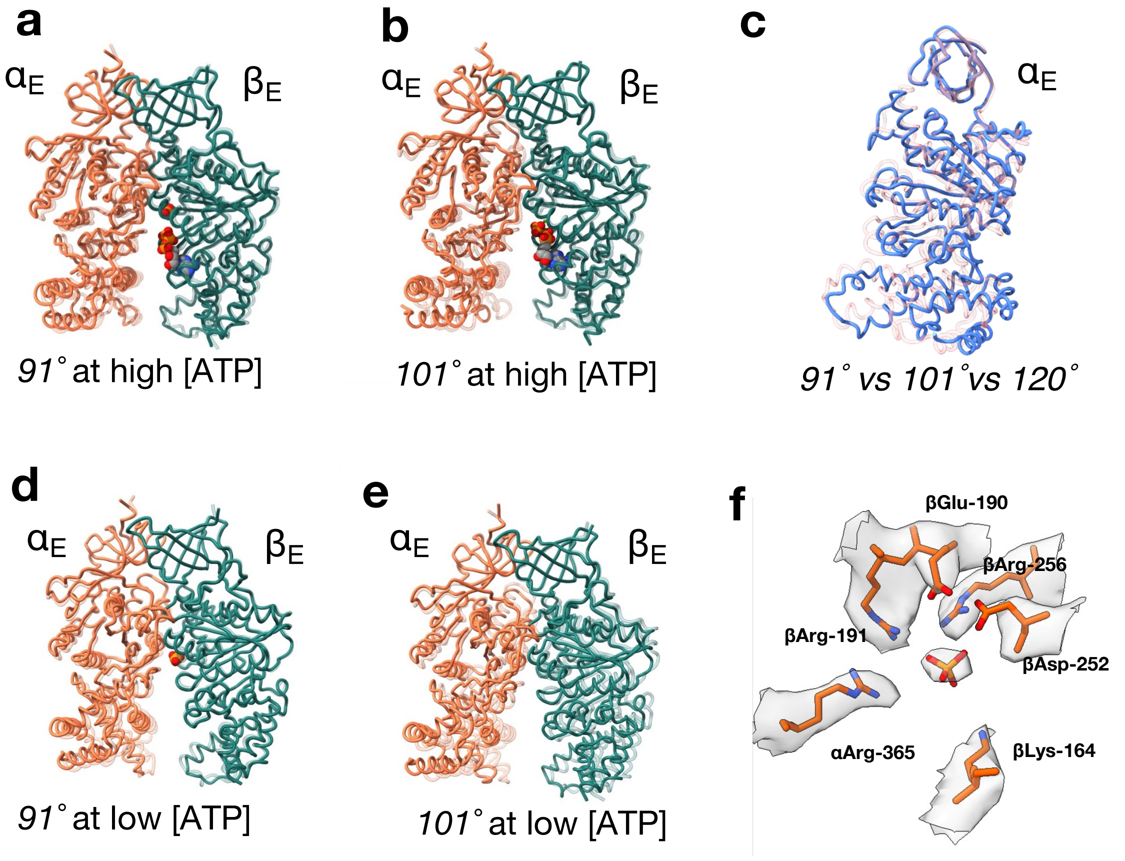


**Extended Data Figure 6.** Conformational changes of $\alpha_E\beta_E$ during the 40° rotation step. **a** Transparent $\alpha_E\beta_E$ in *post-hyd* is superimposed with $\alpha_E\beta_E$ in *91°* at high [ATP]. $\alpha_E$ and $\beta_E$ are colored orange and green respectively. Bound ADP and Pi are represented by spheres. **b** Transparent $\alpha_E\beta_E$ in *91°* is superimposed with $\alpha_E\beta_E$ in *101°* at high [ATP]. **c** Superimposition of $\alpha_E$ in *120°(light blue chain)* with $\alpha_E$ in *101°(pink chain)* and $\alpha_E$ in *91°(transparent pink chain)*. **d** Transparent $\alpha_E\beta_E$ in *post-hyd* superimposed with $\alpha_E\beta_E$ in *91°* at low [ATP]. **e** Transparent $\alpha_E\beta_E$ in *91°* superimposed with $\alpha_E\beta_E$ in *101°* at low [ATP]. **f** Bound Pi and coordinated amino acid residues in $\alpha_E\beta_E$ of *post-hyd* with the density map.

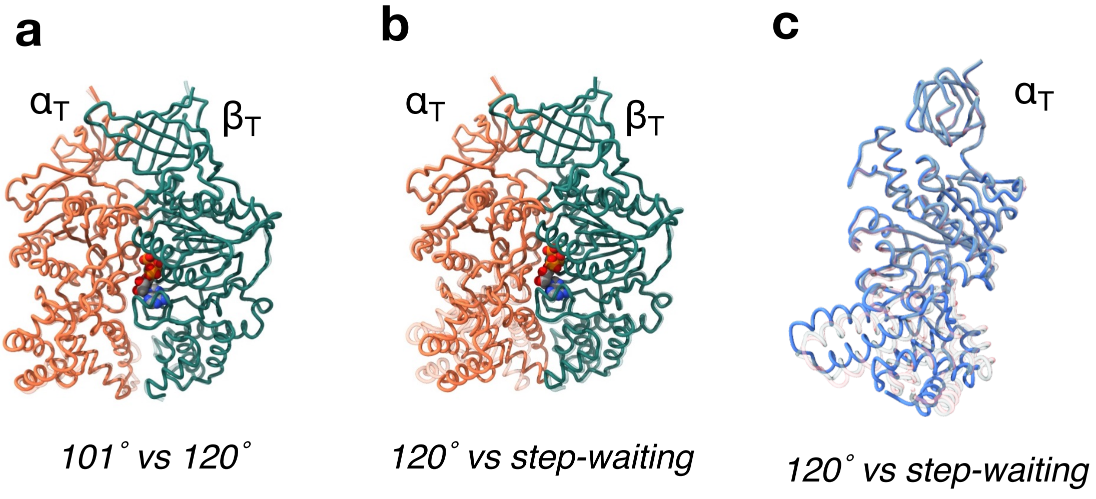


**Extended Data Figure 7.** Conformational changes of $\alpha_T$ between *120°* and *step-waiting*. **a** Transparent $\alpha_T\beta_T$ in *101°* is superimposed with $\alpha_T\beta_T$ in *120°*. The α and β subunits are colored and green, respectively. Bound ATP is represented by spheres. **b** Transparent $\alpha_T\beta_T$ in *120°* is superimposed with $\alpha_T\beta_T$ in *step-waiting*. **c** Superimposition of $\alpha_T$ subunit in *step-waiting (light blue chain)* with α subunit in *120° (transparent pink chain)*.

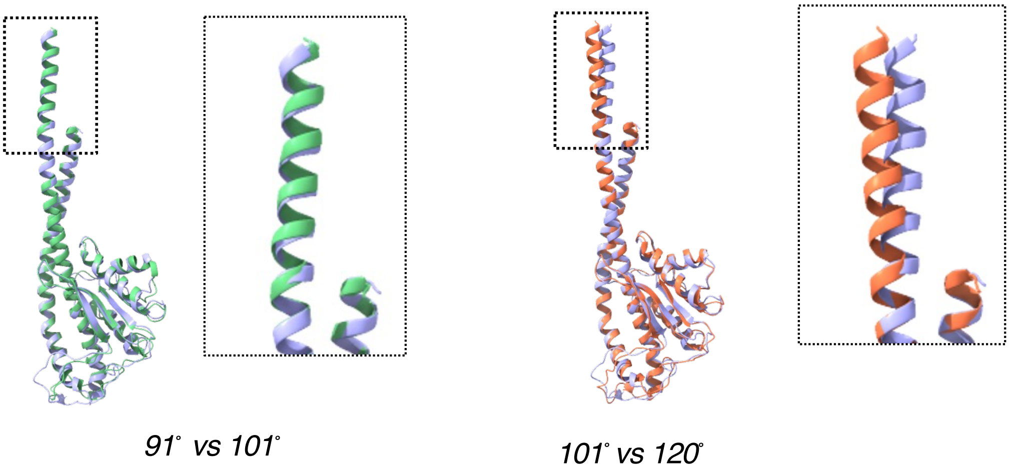


**Extended Data Figure 8.** Conformational changes of γ subunit between *91°*(green) and *101°*(blue) (**a**), and *101°* and *120°*(orange) (**b**). The magnified views of the C termini helices are shown in the right panels, respectively.

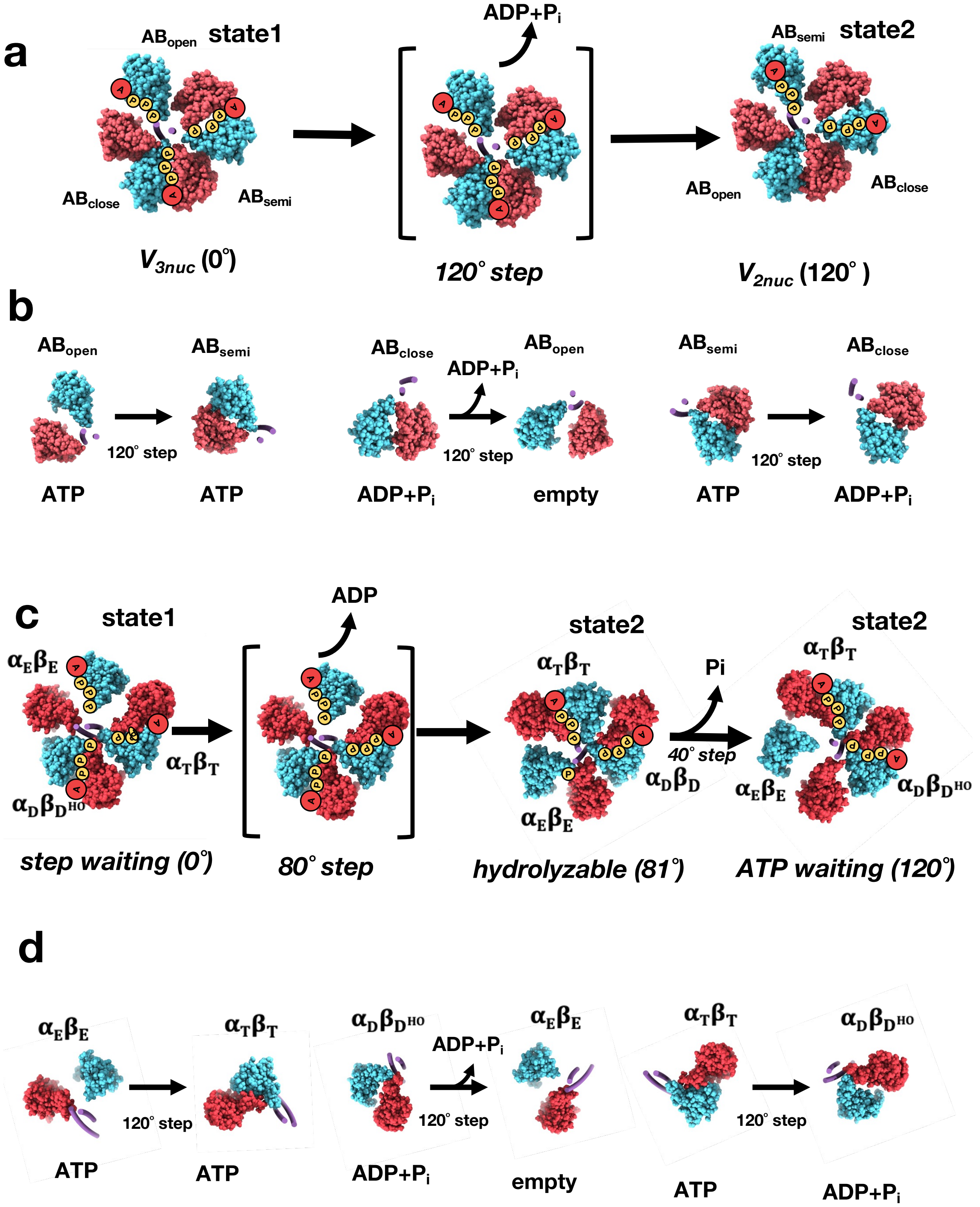


**Extended Data Figure 9. a** Structural change of the $V_1$ domain in V/A-ATPase during a complete 120˚ rotation. $V_{3nuc}$, generated by binding of ATP to $V_{2nuc}$ in state 1, initiates the 120˚ rotation, resulting in $V_{2nuc}$ in state 2. **b** Three catalytic events occur simultaneously at the three catalytic sites; closure of $AB_{open}$ caused by binding of ATP, release of ADP and Pi from $AB_{closed}$ by an opening motion of $AB_{closed}$, and hydrolysis of ATP in $AB_{semi}$, coupled with the 120° rotation of the central DF stalk. **c** Rotation scheme for $F_1$ domain in $F_oF_1$ during the 120° step. **d** During the 120° rotation of the $\gamma$ subunit, three catalytic events also occur simultaneously at the three catalytic sites; closure of $\alpha_E\beta_E$ caused by binding of ATP, release of ADP and Pi from $\alpha_D\beta_D$ by an opening motion of $\alpha_D\beta_D$, and hydrolysis of ATP in $\alpha_T\beta_T$.

**Extended Data Table 1** RMSD values for $F_1$ domain of each s. The $F_1$ domains were superimposed on the β/10-80 a.a. and α/30-90 a.a., then the values for the back bone of $F_1$ domain were calculated using UCSF Chimera software. The lowest value in each row is indicated in bold.

| low [ATP] \ high [ATP] | *hydrolysable* | *post-hyd* | *91°* | *101°* | *120°* | *step-waiting* |
|---|---|---|---|---|---|---|
| *hydrolysable* | **0.79** | 1.25 | 2.00 | 2.71 | 2.77 | 2.71 |
| *post-hyd* | 1.01 | **1.08** | 1.85 | 2.43 | 2.48 | 3.22 |
| *91°* | 1.57 | 1.41 | **1.68** | 1.99 | 2.01 | 2.84 |
| *101°* | / | / | / | / | / | / |
| *120°* | 2.69 | 2.55 | 2.36 | 1.38 | **0.77** | 1.82 |
| *ATP-waiting* | 3.25 | 3.14 | 2.79 | 1.72 | 1.53 | **0.66** |

**Extended Data Table 2a** Cryo-EM data collection, refinement and validation statistics for $F_oF_1$ at high [ATP]

| | *hydrolyzable* | *post-hyd* | *91°* | *101°* | *120°* | *step waiting* |
|---|---|---|---|---|---|---|
| EMDB ID | 34748 | 34749 | 34750 | 34751 | 134752 | 34753 |
| PDB ID | 8HH1 | 8HH2 | 8HH3 | 8HH4 | 8HH5 | 8HH6 |
| **Data collection and processing** | | | | | | |
| Magnification | 81,000 | 81,000 | 81,000 | 81,000 | 81,000 | 81,000 |
| Voltage(kV) | 300 | 300 | 300 | 300 | 300 | 300 |
| Microscope | Titan Krios | Titan Krios | Titan Krios | Titan Krios | Titan Krios | Titan Krios |
| Total dose ($e^-$/Å$^2$) | 60 | 60 | 60 | 60 | 60 | 60 |
| Pixel size(Å/pix) | 0.88 | 0.88 | 0.88 | 0.88 | 0.88 | 0.88 |
| Defocus range(μm) | -0.8 to -2.0 | -0.8 to -2.0 | -0.8 to -2.0 | -0.8 to -2.0 | -0.8 to -2.0 | -0.8 to -2.0 |
| symmetry imposed | C1 | C1 | C1 | C1 | C1 | C1 |
| Initial particle | 1,118,093 | 1,118,093 | 1,118,093 | 1,118,093 | 1,118,093 | 1,118,093 |
| Final Particle | 36,916 | 19,470 | 6,516 | 15,893 | 14,694 | 26,536 |
| Map resolution(Å) | 2.9 | 3.0 | 4.3 | 3.1 | 2.9 | 2.9 |
| FSC threshold | 0.143 | 0.143 | 0.143 | 0.143 | 0.143 | 0.143 |
| **Refinement** | | | | | | |
| Initial model used | This study | This study | This study | This study | This study | This study |
| Model resolution | 2.9 | 3.1 | 4.3 | 3.3 | 3.2 | 3.2 |
| FSC threshold | 0.5 | 0.5 | 0.5 | 0.5 | 0.5 | 0.5 |
| **Model composition** | | | | | | |
| Nonhydrogen atoms | 24280 | 24282 | 24288 | 24284 | 24284 | 24284 |
| Protein residues | 3129 | 3129 | 3129 | 3129 | 3129 | 3129 |
| Ligands | 5MG,6ATP, ,1$PO_4$ | 5MG,5ATP, 1ADP,2$PO_4$ | 5MG,5ATP, 1ADP,2$PO_4$ | 6MG,5ATP, 1ADP,1$PO_4$ | 6MG,5ATP, 1ADP,1$PO_4$ | 6MG,5ATP, 1ADP,1$PO_4$ |
| **R.m.s deviations** | | | | | | |
| Bond length (Å) | 0.002 | 0.004 | 0.003 | 0.004 | 0.003 | 0.002 |
| Bond Angles (°) | 0.54 | 0.647 | 0.617 | 0.646 | 0.609 | 0.585 |
| **Validation** | | | | | | |
| MolProbity score | 1.19 | 1.22 | 1.92 | 1.43 | 1.35 | 1.32 |
| EMRinger score | 3.96 | 3.2 | 1.03 | 3.01 | 3.47 | 3.28 |
| Clashscore | 4.1 | 4.06 | 10.3 | 4.12 | 5.62 | 4.98 |
| Rotamer outlier (%) | 0 | 0.51 | 0.12 | 0.04 | 0 | 0 |
| CaBALM outlier (%) | 1.61 | 1.84 | 2.87 | 2.26 | 1.68 | 1.84 |
| **Ramachandran plot** | | | | | | |
| Favored (%) | 98.11 | 97.88 | 94.35 | 96.44 | 97.78 | 97.72 |
| Allowed (%) | 1.93 | 2.05 | 5.65 | 3.53 | 2.18 | 2.28 |
| Disallowed (%) | 0 | 0.06 | 0 | 0.03 | 0 | 0 |

**Extended Data Table 2b** Cryo-EM data collection, refinement and validation statistics for $F_oF_1$ at low [ATP].

| | *hydrolyzable* | *post-hyd* | *Post-hyd'* | *91°* | *120°* | *step waiting* |
|---|---|---|---|---|---|---|
| EMDB ID | 34754 | 34755 | 34760 | 34756 | 34757 | 34758 |
| PDB ID | 8HH7 | 8HH8 | 8HHC | 8HH9 | 8HHA | 8HHB |
| **Data collection and processing** | | | | | | |
| Magnification | 81,000 | 81,000 | 81,000 | 81,000 | 81,000 | 81,000 |
| Voltage(kV) | 300 | 300 | 300 | 300 | 300 | 300 |
| Microscope | Titan Krios | Titan Krios | Titan Krios | Titan Krios | Titan Krios | Titan Krios |
| Total dose ($e^-$/Å$^2$) | 60 | 60 | 60 | 60 | 60 | 60 |
| Pixel size(Å/pix) | 0.88 | 0.88 | 0.88 | 0.88 | 0.88 | 0.88 |
| Defocus range(μm) | -0.8 to -2.0 | -0.8 to -2.0 | -0.8 to -2.0 | -0.8 to -2.0 | -0.8 to -2.0 | -0.8 to -2.0 |
| symmetry imposed | C1 | C1 | C1 | C1 | C1 | C1 |
| Initial particle | 2,654,860 | 2,654,860 | 2,654,860 | 2,654,860 | 2,654,860 | 2,654,860 |
| Final Particle | 119,152 | 82,415 | 26,578 | 16,437 | 13,133 | 7,818 |
| Map resolution(Å) | 2.5 | 2.8 | 3.3 | 3.6 | 3.4 | 3.5 |
| FSC threshold | 0.143 | 0.143 | 0.143 | 0.143 | 0.143 | 0.143 |
| **Refinement** | | | | | | |
| Initial model used | This study | This study | This study | This study | This study | This study |
| Model resolution | 2.7 | 3 | 3.4 | 3.6 | 3.5 | 3.5 |
| FSC threshold | 0.5 | 0.5 | 0.5 | 0.5 | 0.5 | 0.5 |
| **Model composition** | | | | | | |
| Nonhydrogen atoms | 24280 | 24282 | 24288 | 24284 | 24284 | 24284 |
| Protein residues | 3129 | 3129 | 3129 | 3129 | 3129 | 3129 |
| Ligands | 5MG,6ATP, ,$1PO_4$ | 5MG,5ATP, 1ADP,$2PO_4$ | 5MG,5ATP, 1ADP,$2PO_4$ | 6MG,5ATP, 1ADP,$1PO_4$ | 6MG,5ATP, 1ADP,$1PO_4$ | 6MG,5ATP, 1ADP,$1PO_4$ |
| **R.m.s deviations** | | | | | | |
| Bond length (Å) | 0.002 | 0.004 | 0.003 | 0.004 | 0.003 | 0.002 |
| Bond Angles (°) | 0.54 | 0.647 | 0.617 | 0.646 | 0.609 | 0.585 |
| **Validation** | | | | | | |
| MolProbity score | 1.19 | 1.22 | 1.92 | 1.43 | 1.35 | 1.32 |
| EMRinger score | 3.96 | 3.2 | 1.03 | 3.01 | 3.47 | 3.28 |
| Clashscore | 4.1 | 4.06 | 10.3 | 4.12 | 5.62 | 4.98 |
| Rotamer outlier (%) | 0 | 0.51 | 0.12 | 0.04 | 0 | 0 |
| CaBALM outlier (%) | 1.61 | 1.84 | 2.87 | 2.26 | 1.68 | 1.84 |
| **Ramachandran plot** | | | | | | |
| Favored (%) | 98.11 | 97.88 | 94.35 | 96.44 | 97.78 | 97.72 |
| Allowed (%) | 1.93 | 2.05 | 5.65 | 3.53 | 2.18 | 2.28 |
| Disallowed (%) | 0 | 0.06 | 0 | 0.03 | 0 | 0 |

**Extended Data Table 2c** Cryo-EM data collection, refinement for $F_oF_1$ at high [ATP]

| | *hydrolyzable* | | | *post-hyd* | | | *90°* | | |
|---|---|---|---|---|---|---|---|---|---|
| | state1 | state2 | state3 | state1 | state2 | state3 | state1 | state2 | state3 |
| EMDB ID | 34759 | 34761 | 34763 | 34762 | 34764 | 34765 | 34766 | 34767 | 34770 |
| **Data collection and processing** | | | | | | | | | |
| Magnification | 81,000 | 81,000 | 81,000 | 81,000 | 81,000 | 81,000 | 81,000 | 81,000 | 81,000 |
| Microscope | Titan Krios | Titan Krios | Titan Krios | Titan Krios | Titan Krios | Titan Krios | Titan Krios | Titan Krios | Titan Krios |
| Voltage(kV) | 300 | 300 | 300 | 300 | 300 | 300 | 300 | 300 | 300 |
| Total dose ($e^-$/Å$^2$) | 60 | 60 | 60 | 60 | 60 | 60 | 60 | 60 | 60 |
| Pixel size(Å/pix) | 0.88 | 0.88 | 0.88 | 0.88 | 0.88 | 0.88 | 0.88 | 0.88 | 0.88 |
| Defocus range(μm) | -0.8 to -2.0 | -0.8 to -2.0 | -0.8 to -2.0 | -0.8 to -2.0 | -0.8 to -2.0 | -0.8 to -2.0 | -0.8 to -2.0 | -0.8 to -2.0 | -0.8 to -2.0 |
| symmetry imposed | C1 | C1 | C1 | C1 | C1 | C1 | C1 | C1 | C1 |
| Initial particle | 1,381,269 | 1,381,269 | 1,381,269 | 1,381,269 | 1,381,269 | 1,381,269 | 1,381,269 | 1,381,269 | 1,381,269 |
| Final Particle | 20,667 | 7,951 | 7,916 | 10,472 | 4,621 | 4,041 | 3,432 | 2,141 | 937 |
| Map resolution(Å) | 3.0 | 3.3 | 3.3 | 3.4 | 3.8 | 3.8 | 6.7 | 8.0 | 8.8 |
| FSC threshold | 0.143 | 0.143 | 0.143 | 0.143 | 0.143 | 0.143 | 0.143 | 0.143 | 0.143 |

| | *100°* | | | *120°* | | | *step waiting* | | |
|---|---|---|---|---|---|---|---|---|---|
| | state1 | state2 | state3 | state1 | state2 | state3 | state1 | state2 | state3 |
| EMDB ID | 34768 | 34769 | 34770 | 34771 | 34772 | 34773 | 34774 | 34775 | 34776 |
| **Data collection and processing** | | | | | | | | | |
| Magnification | 81,000 | 81,000 | 81,000 | 81,000 | 81,000 | 81,000 | 81,000 | 81,000 | 81,000 |
| Microscope | Titan Krios | Titan Krios | Titan Krios | Titan Krios | Titan Krios | Titan Krios | Titan Krios | Titan Krios | Titan Krios |
| Voltage(kV) | 300 | 300 | 300 | 300 | 300 | 300 | 300 | 300 | 300 |
| Total dose ($e^-$/Å$^2$) | 60 | 60 | 60 | 60 | 60 | 60 | 60 | 60 | 60 |
| Pixel size(Å/pix) | 0.88 | 0.88 | 0.88 | 0.88 | 0.88 | 0.88 | 0.88 | 0.88 | 0.88 |
| Defocus range(μm) | -0.8 to -2.0 | -0.8 to -2.0 | -0.8 to -2.0 | -0.8 to -2.0 | -0.8 to -2.0 | -0.8 to -2.0 | -0.8 to -2.0 | -0.8 to -2.0 | -0.8 to -2.0 |
| symmetry imposed | C1 | C1 | C1 | C1 | C1 | C1 | C1 | C1 | C1 |
| Initial particle | 1,381,269 | 1,381,269 | 1,381,269 | 1,381,269 | 1,381,269 | 1,381,269 | 1,381,269 | 1,381,269 | 1,381,269 |
| Final Particle | 7,453 | 3,605 | 4,835 | 5,866 | 3,694 | 5,048 | 13,351 | 5,600 | 7,267 |
| Map resolution(Å) | 3.6 | 4.1 | 3.8 | 3.6 | 3.9 | 3.8 | 3.3 | 3.8 | 3.6 |
| FSC threshold | 0.143 | 0.143 | 0.143 | 0.143 | 0.143 | 0.143 | 0.143 | 0.143 | 0.143 |

**Extended Data Table 2d** Cryo-EM data collection, refinement for $F_oF_1$ at low [ATP].

| | *hydrolyzable* | | | *post-hyd* | | *post-hyd'* | |
|---|---|---|---|---|---|---|---|
| | state1 | state2 | state3 | state1 | state2 | state1 | state2 |
| EMDB ID | 34777 | 34779 | 34780 | 34778 | 34781 | 34782 | 23783 |
| **Data collection and processing** | | | | | | | |
| Magnification | 81,000 | 81,000 | 81,000 | 81,000 | 81,000 | 81,000 | 81,000 |
| Microscope | Titan Krios | Titan Krios | Titan Krios | Titan Krios | Titan Krios | Titan Krios | Titan Krios |
| oltage(kV) | 300 | 300 | 300 | 300 | 300 | 300 | 300 |
| Total dose ($e^-$/Å$^2$) | 60 | 60 | 60 | 60 | 60 | 60 | 60 |
| Pixel size(Å/pix) | 0.88 | 0.88 | 0.88 | 0.88 | 0.88 | 0.88 | 0.88 |
| Defocus range(μm) | -0.8 to -2.0 | -0.8 to -2.0 | -0.8 to -2.0 | -0.8 to -2.0 | -0.8 to -2.0 | -0.8 to -2.0 | -0.8 to -2.0 |
| symmetry imposed | C1 | C1 | C1 | C1 | C1 | C1 | C1 |
| Initial particle | 2,654,860 | 2,654,860 | 2,654,860 | 2,654,860 | 2,654,860 | 2,654,860 | 2,654,860 |
| Final Particle | 74,851 | 48, 930 | 38,282 | 42,326 | 33,731 | 11,438 | 14,975 |
| Map resolution(Å) | 2.9 | 3.0 | 3.1 | 3.1 | 3.2 | 3.7 | 3.8 |
| FSC threshold | 0.143 | 0.143 | 0.143 | 0.143 | 0.143 | 0.143 | 0.143 |

| | *91°* | | *120°* | | *step waiting* | | |
|---|---|---|---|---|---|---|---|
| | state1 | state3 | state1 | state2 | state1 | state2 | state3 |
| EMDB ID | 34784 | 34785 | 6,178 | 3,669 | 34788 | 34,789 | 34,790 |
| **Data collection and processing** | | | | | | | |
| Magnification | 81,000 | 81,000 | 81,000 | 81,000 | 81,000 | 81,000 | 81,000 |
| Microscope | Titan Krios | Titan Krios | Titan Krios | Titan Krios | Titan Krios | Titan Krios | Titan Krios |
| Voltage(kV) | 300 | 300 | 300 | 300 | 300 | 300 | 300 |
| Total dose ($e^-$/Å$^2$) | 60 | 60 | 60 | 60 | 60 | 60 | 60 |
| Pixel size(Å/pix) | 0.88 | 0.88 | 0.88 | 0.88 | 0.88 | 0.88 | 0.88 |
| Defocus range(μm) | -0.8 to -2.0 | -0.8 to -2.0 | -0.8 to -2.0 | -0.8 to -2.0 | -0.8 to -2.0 | -0.8 to -2.0 | -0.8 to -2.0 |
| symmetry imposed | C1 | C1 | C1 | C1 | C1 | C1 | C1 |
| Initial particle | 2,654,860 | 2,654,860 | 2,654,860 | 2,654,860 | 2,654,860 | 2,654,860 | 2,654,860 |
| Final Particle | 7,450 | 7,076 | 6,178 | 3,669 | 10,739 | 5,923 | 2,512 |
| Map resolution(Å) | 4.2 | 4.2 | 3.9 | 4.2 | 3.9 | 4.2 | 7.0 |
| FSC threshold | 0.143 | 0.143 | 0.143 | 0.143 | 0.143 | 0.143 | 0.143 |

**Supplementary movie 1. Structural changes in F1 domain during 120° step.**
